# Partitioning stable and unstable expression level variation in cell populations: a theoretical framework with application to the T cell receptor

**DOI:** 10.1101/527663

**Authors:** Thiago S. Guzella, Vasco M. Barreto, Jorge Carneiro

## Abstract

Phenotypic variation in the copy number of gene products expressed by cells or tissues has been the focus of intense investigation. To what extent the observed differences in cellular expression levels are persistent or transient is an intriguing question. Here, we develop a quantitative framework that resolves the expression variation into stable and unstable components. The difference between the expression means in two cohorts isolated from any cell population is shown to converge to an asymptotic value, with a characteristic time, *τ*_*T*_, that measures the timescale of the unstable dynamics. The asymptotic difference in the means, relative to the initial value, measures the stable proportion of the original population variance 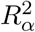. Empowered by this insight, we analysed the T-cell receptor (TCR) expression variation in CD4 T cells. About 70% of TCR expression variance is stable in a diverse polyclonal population, while over 80% of the variance in an isogenic TCR transgenic population is volatile. In both populations the TCR levels fluctuate with a characteristic time of 32 hours. This systematic characterisation of the expression variation dynamics, relying on time series of cohorts’ means, can be combined with technologies that measure gene or protein expression in single cells or in bulk.

## 1 Introduction

The phenotypic variation among organisms or cells is a theme of growing importance in biology. Macroscopic phenotypes, such as body structures or physiologic responses, have been studied for ages, but one phenotype particularly suitable for quantification that has received attention in the last decades is the amount of specific mRNAs and proteins expressed by single cells. Advances in genomics have allowed the analysis of genetic contributions to variation in gene expression, in terms of so-called expression quantitative trait loci (eQTL) [1, 2]. In this case, expression levels, typically assessed via mRNA levels, are treated as quantitative traits, and one is interested in the specific loci underlying variation in expression levels among different individuals. The increasing availability of single-cell resolution genomics, proteomics and metabolomics technologies has enabled molecular biologists to analyse cell lineages and tissues showing that what were previously perceived as homogeneous cell populations are in fact a complex mixture of often transient and interchangeable cellular types and cellular states (see discussion in [3]). In parallel to these studies linking phenotypes to genotype, the literature on stochastic gene expression [4, 5, 6, 7, 8], reviewed in [9], has brought to light the variation in expression levels in isogenic cells, even when these are in the same cellular state and in the same environment. The variation is typically attributed to the “noise” resulting from the small copy number of molecules involved in the process.

Several studies addressed the fluctuation dynamics of gene expression levels [10, 11] revealing a complex picture of the variation in isogenic cell populations. The fluctuation timescales range from hours [7, 12], to days [13, 14, 15] or weeks [16, 17, 18], depending on the cells and on the degree of multimodality of the expression distribution under study. The distinct timescales can be associated with the different mechanisms that may cause the variation in the expression levels of a molecular component of interest in some cell population. However, most quantitative approaches developed up to this date have focused on noise in gene expression as the predominant mechanism explaining the variation observed (for example, [6, 7, 19, 20, 21, 22, 23, 24]). It remains unclear to which degree less volatile dynamic processes or even persistent differences contribute to the observed variation in a isogenic cell population. This is particularly relevant in the case of cells from multicellular organisms, due to the robust epigenetic processes that underlie differentiation stages, cell lineages or cell states, but also the intraclonal structure of apparently homogenous populations [25, 26].

A case study of particular interest is the expression of Sca1 in a hematopoietic cell line [16, 17] since it reveals the inherent complexity of variation dynamics and also the difficulties in characterising it experimentally. Chang et al. [16] reported that that biased cohorts of cells tend to restore the histogram of expression levels of Sca1 of the starting population, albeit with very slow dynamics. In principle, complete restoration would be consistent with a lack of stable variants in the population, at least in terms of expression levels of Sca1. However, [17] have shown that, even after 2 weeks, the reconstitution is not fully complete. More importantly, these authors [17] showed that some cells in this population express markers indicative of terminal differentiation, and have limited capacity for cell division. This points to an inherent heterogeneity in the population that may persist in time. An important limitation of these approaches was relying mainly on the juxtaposition of histograms of expression levels in order to compare cell populations, without a rigorous quantification. It is not clear how to analyse such data and because of this the degree to which the original distribution is restored remains uncertain. A quantitative approach that overcomes this impasse is necessary and also important to provide formal concepts on which to ground subsequent studies on the expression levels in cell populations.

Our work lumps the molecular mechanisms regulating expression levels in a cell population into two components, one stable and another unstable. The stable component leads to permanent differences between the expression levels of any two cohorts of cells. The unstable component, on the other hand, represents transient differences in the expression levels of the cohorts, which will eventually vanish in time. Starting from these definitions, a general model is derived to describe protein expression levels in a population with both the stable and unstable components. The relative contribution of the stable component to the expression variation is then defined as a single parameter termed 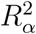. We show theoretically that this parameter can be estimated in an unbiased way by following over time the mean expression in cohorts isolated from the population of interest. This dynamical characterisation of the expression variation is completed by concomitantly estimating the characteristic timescale *τ*_*T*_.

This theoretical result is used to characterise the contributions to variation in expression levels of the T-cell receptor (TCR) in two biologically relevant CD4 T cell populations. The first population, purified from wild type mice, is composed of clones emerging from the process of V(D)J recombination, each carrying genetically distinct TCR loci. The second is a genetically uniform population isolated from Marilyn TCR-transgenic mice, in which all T cells express the same recombined receptor genes [27]. We find that the stable component is the main contribution in the polyclonal population 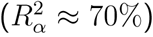, while the unstable component predominates in the Marilyn population 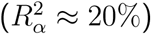. This suggests that genetic heterogeneity contributes to stable differences in TCR expression levels in T cells, but that there are other mechanisms contributing to persistent expression variation in isogenic populations.

## 2. A general model for protein expression levels in a cell population

### 2.1 Partitioning the contributions to variation in expression levels

We assume that any biological cell population, hereafter referred to as full population, is a mixture of sub-populations. Each cell belongs to and remains in one of these sub-populations all the time. Using a mixture model formulation, each sub-population is indexed by *i* = 1, 2, …, *N*, and described by three parameters (*µ*_*i*_, *v*_*i*_, *w*_*i*_): the mean *µ*_*i*_ and variance *v*_*i*_ of expression levels, and the relative frequency *w*_*i*_ of cells in the full population that belong to this sub-population. The latter is given by:

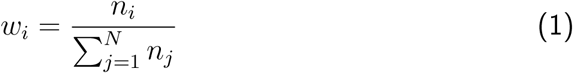

where *n*_*i*_ is the number of cells in the *i*-th sub-population and 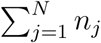 is the total number of cells in the full population. A related approach has been used by Gianola *et al*. [28] to study genetic parameters in the context of the quantitative genetics of mixture characters.

In the limit of large *N*, the parameters (*µ*_*i*_, *v*_*i*_, *w*_*i*_) describing a sub-population are taken as random variables (***µ***, ***v***, ***w***) (see methods for details of the notation used) following a particular multivariate distribution. Then, one can relate the mean *µ*_*F*_ and variance *v*_*F*_ of expression levels of the full population to the properties of the sub-populations, as detailed in section A of the supplementary information. Provided that there is no correlation between the frequencies (***w***) and either the means (***µ***), the squared means (***µ***^2^) and the variances (***v***) of the sub-population, it follows that (supplementary information, section A):

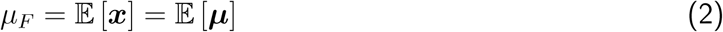

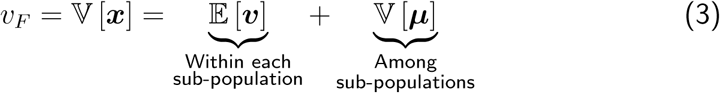

where the subscript *F* is used to highlight that these are properties of the full population. Therefore, under these conditions, the mean of the full population is simply the expected value of the means of the sub-populations (𝔼 [***µ***]), while the variance of the full population is the sum of the variance within each subpopulation (𝔼 [***v***]) and the variance among the sub-populations (variance in the means, 𝕍 [***µ***]).

It is important to highlight that equations 2 and 3 are general and independent of the precise definition of a sub-population. However, the two terms in equation suggest a specific definition, in which only the unstable component is present in each sub-population. In this way, the term of variation within any sub-population 𝔼 [***v***] becomes the contribution of the unstable component to the variance of the full population, while the variation among the means of the sub-populations 𝕍 [***µ***] is the contribution of the stable component. In the next section 2.2, expression levels within each sub-population will be described by a stochastic model, while the different sub-populations will have different means controlled by one of the parameters of this stochastic model.

### 2.2 An explicit model of protein expression in a cell population

#### 2.2.1 Describing variation within a sub-population

The stochastic model of protein expression considered here is based on previous work [29], and is defined by the following two equations:

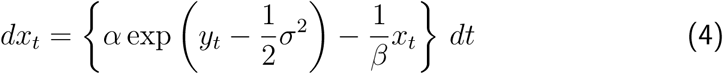

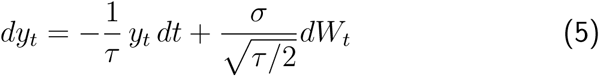

where *x*_*t*_ is the amount of protein expressed at time *t*, and *y*_*t*_ is a stochastic variable following the Ornstein-Uhlenbeck process. In equation 5, *W*_*t*_ is the Wiener process [30]. The parameters for the model are presented in Table 1, along with their respective dimensions.

**Table 1:** Description of the parameters of the stochastic model of protein expression defined by equations 4 and 5

| Parameter | Description | Dimensions |
| --- | --- | --- |
| $\alpha$ | Mean protein production rate | Molecules/Time |
| $\beta$ | Mean lifetime of the protein | Time |
| $\sigma$ | Normalised dispersion of protein production rate | Non-dimensional |
| $\tau$ | Characteristic time of the fluctuations in protein production rate | Time |

The equation governing *dx*_*t*_ has two terms. The first term, 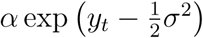, is the rate of production which depends on the stochastic process *y*_*t*_, and the second term, *x*_*t*_/*β*, is the degradation rate following first-order kinetics with mean *β* protein lifetime. A model with a similar overall structure was reported before [31], in which mRNA transcription and degradation have also been explicitly incorporated. Equation 4 can be re-written as:

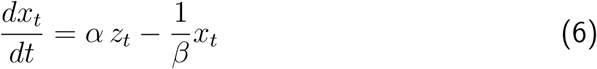

where *z*_*t*_, defined as:

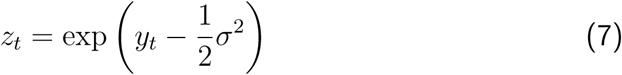

denotes the instantaneous rate of protein production. This rate is normalised, to have unit expected value. All processes governing protein production (promoter transitions, transcription and translation, among others) are lumped together into the average rate *α* and the instantaneous rate given by *z*_*t*_. The representation in equation 6, which highlights the contribution of lumped upstream factors, has been applied before in the analysis of models of stochastic gene expression (for example, [6, 7]). Equation 6 denotes that, in a single cell, the instantaneous rate of protein production is proportional to the instantaneous levels of these lumped upstream factors, and fluctuates as a function of time, with auto-correlation time approximately equal to *τ* [29]. These fluctuations are then propagated downstream, resulting in fluctuations in protein levels, with dynamics dictated by *τ* (through *z*_*t*_) and *β*. For simplicity, protein degradation is assumed to be deterministic, with the same rate 1/*β* for all cells. The temporal evolution of the protein expression levels *x*_*t*_ in two cells with distinct characteristic times *τ* is illustrated in figure 1 (top).

**Figure 1:**
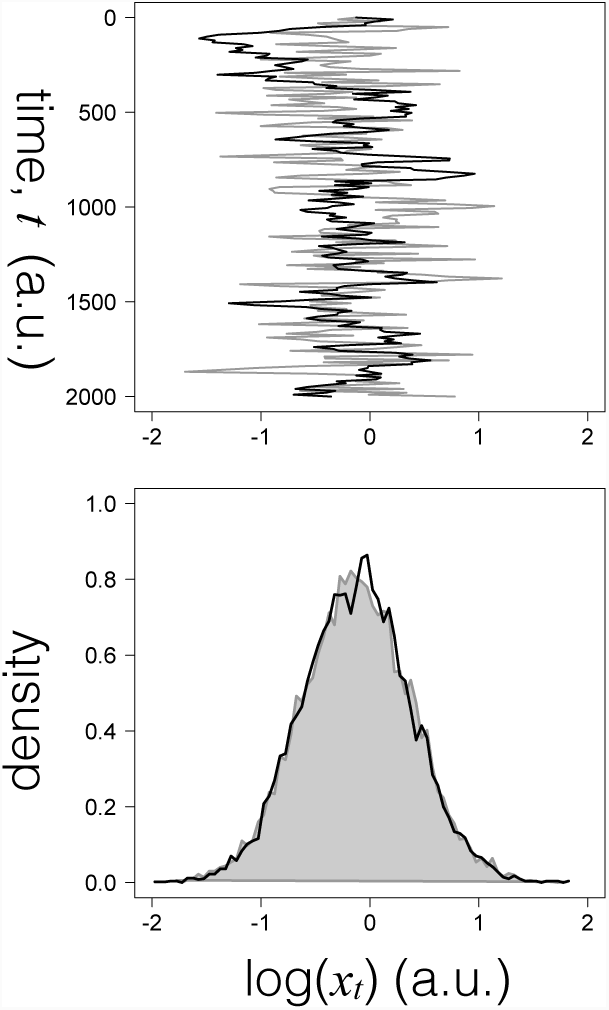
Dynamics of the protein expression levels *x*_*t*_ according to the stochastic model. Top-Time course of the log-transformed variable *x*_*t*_ obtained for two cells which differ in the characteristic time of the fluctuations (*τ* = 10 (grey) and *τ* = 100 (black)). BottomHistograms of the log-transformed protein levels *x*_*t*_ in cell populations with slow and fast dynamics exemplified by the time courses. Each histogram corresponds to 10000 independent realisations of the individual cell model sampled at time *t* = 200*a.u*.. The histograms were normalised by their maximum density. Remaining parameters: *α* = 1., *β* = 1, and *σ* = 0.5.

It follows from equation 7 that:

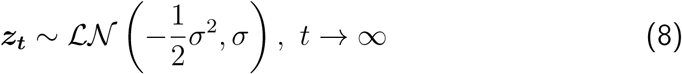

and therefore the stationary rate of protein production follows a lognormal distribution in cells of a sub-population, consistent with a report of lognormal rates of protein expression [32]. Equations 4 and 5 are a simple model that generates, for a wide range of parameter values, a lognormal-like distribution of protein levels (figure 1,bottom), compatible with the widespread observation of the lognormal distribution in cell populations. In this scenario, in terms of the log-transformed protein levels (section B of the supplementary information), the mean and variance of a stationary sub-population are given by equations 9 and 10, respectively:

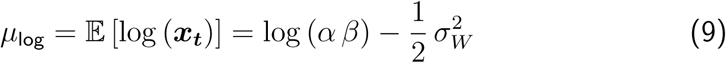

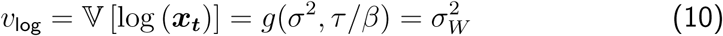

where the subscript *W* will be used hereafter to denote that the variation is due to the stochastic process influencing the instantaneous rate of protein production. In equation 10, *g*(*·, ·*) is an arbitrary function, which can be estimated via simulation.

#### 2.2.2 Combining variation within and among sub-populations

As formulated in section 2.1, the stable component arises due to variation in the means of the sub-populations. Therefore, we assume that parameter *α* in equation 4 is distributed in the full population, becoming a random variable, denoted by ***α***. Consequently, each sub-population is described by one value of *α*, resulting in different average rates of production, and hence different mean expression levels.

For simplicity, we consider the case that ***α*** *∼ℒ 𝒩 (µ*_*α*_, *σ*_*α*_). For the *i*-th sub-population, with parameter *α* _*i*_, the mean and variance follow from equations 9 and 10:

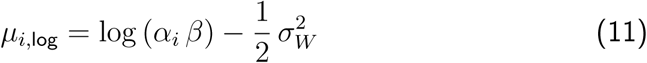

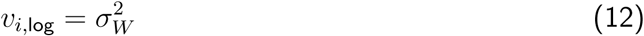

where 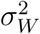 is assumed to be the same for all sub-populations. In terms of log-transformed values, plugging equations 9 and 10 into equation 3, one obtains the variance of the full population:

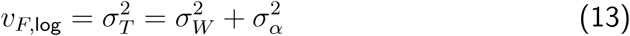

An important property of equation 13, which is based on log-transformed values, is that the parameters that represent the variances due to the stable and unstable components (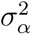 and 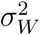, respectively) remain separate. This is a key feature, greatly simplifying the process of analysis and inference throughout this work. As detailed in the supplementary information (section C), the equivalent of equation 13 considering protein levels without any transformation has an additional term, dependent on 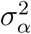 and 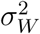. This additional term arises since the variance of each sub-population in this case depends on the value of *α*. Therefore, we consider, from this point on, the analysis based on log-transformed values only.

## 3 Isolating cells to quantify the contributions to the variation in a cell population

### 3.1 Defining the relative contribution of the stable component

The previous section 2.2.2 showed that the variance of log-transformed expression levels of the full population is simply the sum of variances due to the stable and unstable components (equation 13). In this context, in analogy with the *R*^2^ quantification of the variance explained by a linear regression model, we define 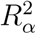 as:

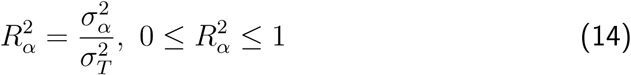

to denote the proportion of the observed variance that is explained by the stable component.

Hence, 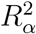 formalizes and quantifies the relative contribution of the stable component to the total variance of the full population, reducing the problem of quantifying the contributions to the estimation of a single parameter. In the case of 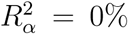, variation in expression levels arises exclusively due to the unstable component; conversely, the stable component explains all the observed variation if 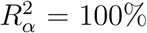. Finally, in the intermediate case 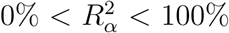, a combination of the two components is at play.

### 3.2 Estimating the relative contribution of the stable component by isolating cells

After defining 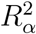, a setup for its estimation is considered. Since the original population is assumed to be heterogeneous, being composed of several subpopulations, a natural approach for estimation is to isolate a cohort of cells and to follow the temporal evolution of some property of this cohort. The isolation of cells according to the expression levels of some protein has been described in previous experimental works [16, 10, 14, 17, 18], usually employing fluorescence-activated cell sorting (FACS). To simplify the presentation, it is assumed that the property is always quantified based on a sufficiently large number of cells, such that sampling effects are negligible.

Hereafter, a time reference *t* is defined beginning from the instant of isolation in a hypothetical experiment. Let an isolated cell cohort correspond to cells between percentiles *p*_1_ and *p*_2_ of expression levels of the original population. Without loss of generality, it is assumed hereafter that *p*_1_ *< p*_2_. Therefore, the two percentiles should satisfy 0% ≤ *p*_1_ *< p*_2_ *<* 100% or 0% *< p*_1_ *< p*_2_ ≤ 100%. This ensures that at least one of the isolated cohorts to be used for inference is not identical to the original population at time *t* = 0. Hence, isolating cells corresponds, indirectly, to selecting some of the sub-populations, if any compose compose the original population. Upon isolation, the expression levels of cells in a given sub-population will relax to the stationary distribution of that subpopulation. Therefore, at the level of the isolated cohorts being tracked, changes in the property of expression levels are related to the dynamics of the unstable component, as expression levels of the sub-populations that have been isolated relax to their stationary values. The time for this relaxation to take place will be hereafter referred to as the characteristic time of the variation.

In a given experiment, three outcomes are possible (figure 2). If only the unstable component is present 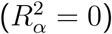, after waiting a sufficiently long amount of time, the distribution of protein expression in the isolated cohort will converge to that of the original population (figure 2, top). In contrast, if the observed variation is explained by the stable component only 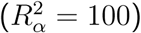, the distribution of the isolated cohort will not change as a function of time, remaining identical to that just after being isolated; it will always differ from that of the original population (figure 2, middle). Finally, if both the stable and unstable components are present in the original population 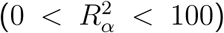, the isolated cohort will evolve in time, but without ever restoring the distribution of the original population (figure 2, bottom).

**Figure 2:**
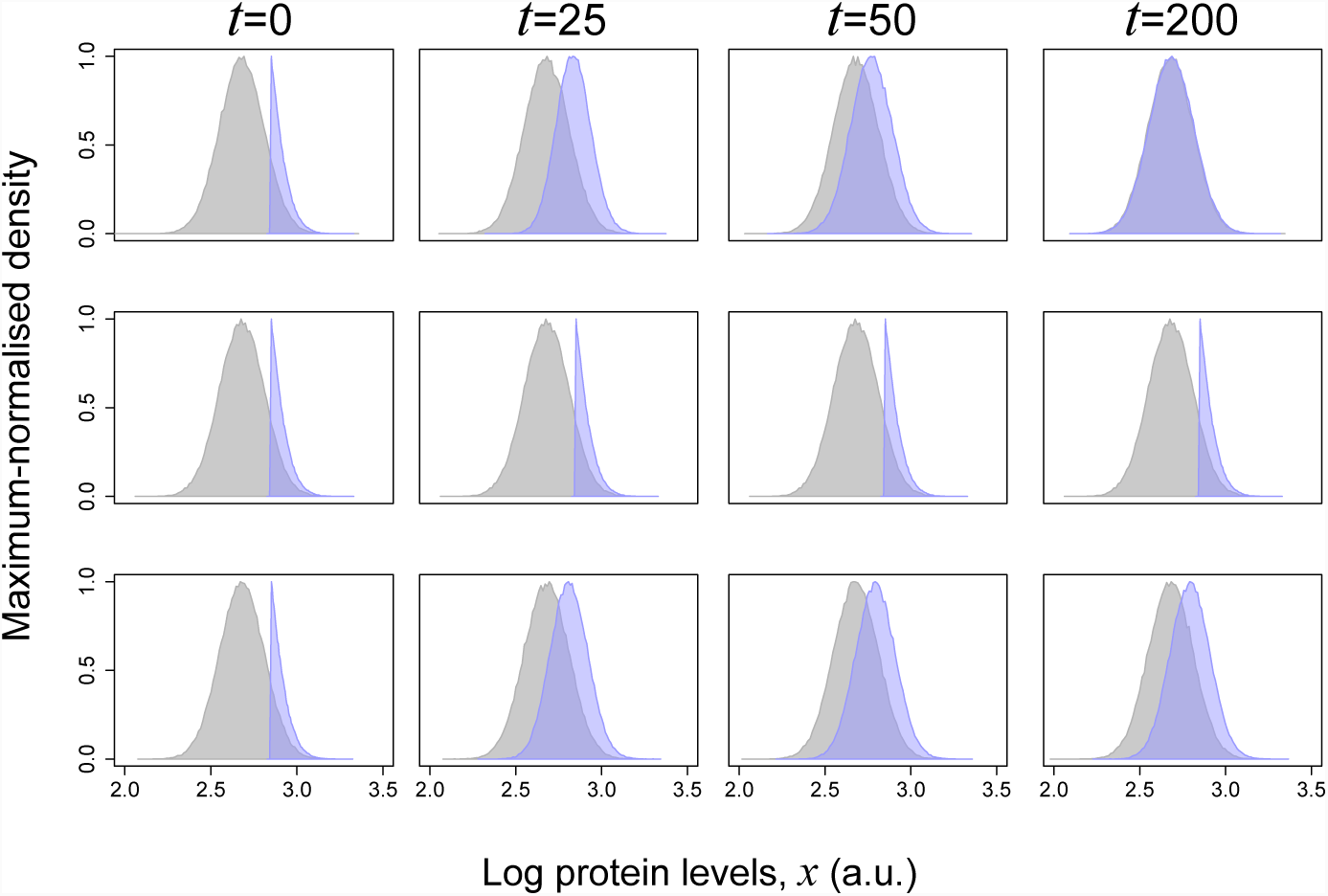
Simulation of the possible results obtained when a cohort of high expressor cells is isolated from a full population and followed in time. The graphs are histograms of the values of the expression levels variable *x* at the indicated times in 10000 independent realisations of the model for three values of 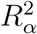 (0.0 (top), 1.0 (middle) and 0.25 (bottom)) simulating an isolated cohort of higher expressors (blue) or the full population from which the cohort was isolated (gray).

The key question now is what properties of the isolated cohort can be used to infer 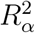. The next section shows that 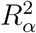 can be accurately inferred from the dynamics of the means of the cohorts and examines the choice of a specific approach for isolation in term of the percentiles *p*_1_ and *p*_2_. The additional features that can be extracted from the variance of the isolated cohort are addressed in supplemental information section E.

## 4 Estimating the relative contribution of the stable component

This section uses simulation to identify which property of the isolated cohorts can lead to a good estimate of 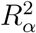, when followed in time. In the simulations, protein expression levels are described by the model derived in section 2, neglecting cell division for simplicity. Since all derivations are based on equation 13, the analysis herein relies on log-transformed values of protein levels.

The isolated cohort considered at first for inference here is composed of the 10% of cells with the highest (respectively lowest) expression levels in the original population hereafter referred to as “high expressors” (respectively “low expressors”). Following the notation of section 3.2, we have *p*_1_ = 90% and *p*_2_ = 100% (respectively, *p*_1_ = 0% and *p*_2_ = 10%). The choice of 10% is arbitrary, and is deemed to represent, at least in principle, a good compromise between resolution and number of cells obtained. Moreover, a random sample of the original population will serve as reference.

We first address how the dynamics of the mean of log-transformed protein levels in isolated cohorts, shown in figure 3 (left) for the high expressors. Briefly, the mean of an isolated cohort will evolve smoothly until it reaches an asymptotic limit.

**Figure 3:**
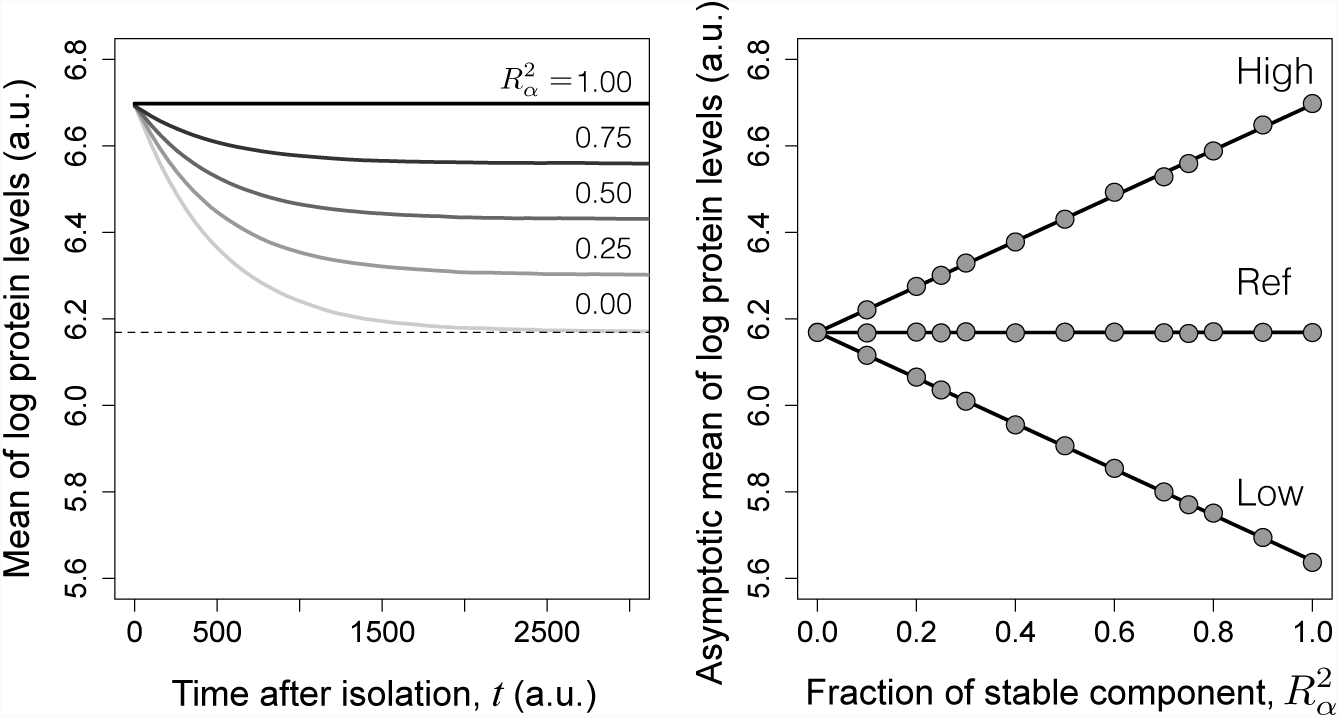
Simulation of transient dynamics and asymptotic limits of the mean protein expression in isolated cohorts. a) Dynamics of mean log protein expression levels of “high expressors” after isolation as 10% of original populations with different values of 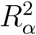, but constant 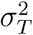. b) Asymptotic mean of 10% low expressors, 10% high expressors and reference population as a function of 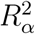 in the simulations. The symbols represent simulation results, while the lines represent the best-fit of a straight line. Remaining parameter values: *τ* = 500, *β* = 5 and *σ*_*T*_ = 0.3.

It turns out that the asymptotic mean of log protein levels of an isolated cohort is a linear function of the 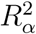 of the original population from which it was obtained, as illustrated in figure 3(right) in the cases of the isolation of the 10% high and low expressors in populations with different 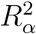. This linear relationship allows one to define a straightforward approach for estimating 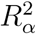. Defining Δ_*A,B*_(*t*) as the difference between the means of log-transformed values of two isolated cohort *A* and *B*, respectively *µ*_*A*_(*t*) and *µ*_*B*_(*t*), at time instant *t*:

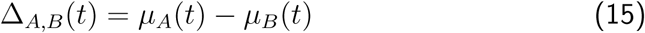

then, 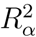 can be estimated via:

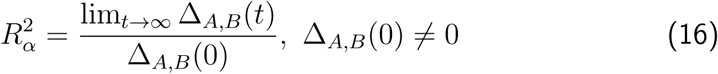

The condition Δ_*A,B*_(0) ≠ 0 for using equation 16 implies that the two isolated cohorts being compared must have different means just after isolation (*t* = 0). From the inequality in equation 14, an additional relationship for Δ_*A,B*_(*t*) holds:

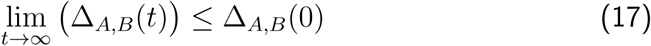

Therefore, the stationary difference between the means of log-transformed expression levels of the isolated cohorts *A* and *B* is expected to be, under the present formulation, lower than or equal to the difference immediately after isolation. Therefore, a key result is that, to estimate 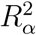, one may simply calculate the ratio between the asymptotic value of difference between the means of logtransformed protein levels in two isolated cohorts relative to its initial value after isolation.

An important consequence for experimental design is that one can improve the resolution in the estimation of 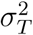 by maximising the value of Δ_*A,B*_(0). For any given percentage of cells to be isolated (the difference *p*_2_ -*p*_1_; see section 3.2), the maximal initial difference is obtained by isolating the extreme high and low expressors. Consequently, the remainder of this work focuses on this case, by always relying on the function Δ_*H,L*_(*t*) for estimation, where H and L denoted respectively the high and low expressors.

Equation 16 has an important advantage from an experimental point of view: the fact that it depends only on the differences between the means of the sorted and reference populations. This is particularly important given that there are typically day-to-day variations in the absolute values read by a flow cytometer, to which equation 16 is robust.

The asymptotic analysis just presented does not allow to consider the dynamics of the expression levels. To address these dynamics we introduce the time-dependent function Ω_*H,L*_(*t*) given by:

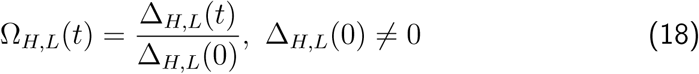

Being based on the means of log-transformed values of two populations that have been isolated, Δ_*H,L*_(*t*) follows an approximately exponential decay (figure 4; see section D of the supplementary material for a rationale). Using the approximation of exponential decay, and defining the effective characteristic time as *τ*_*T*_, Ω_*H,L*_(*t*) takes the following form:

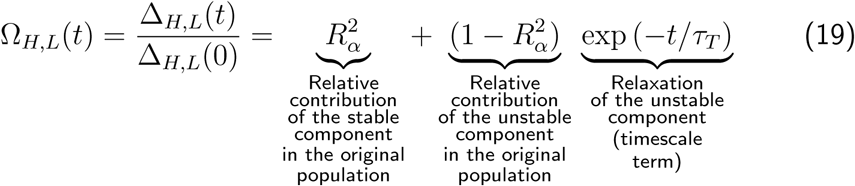

**Figure 4:**
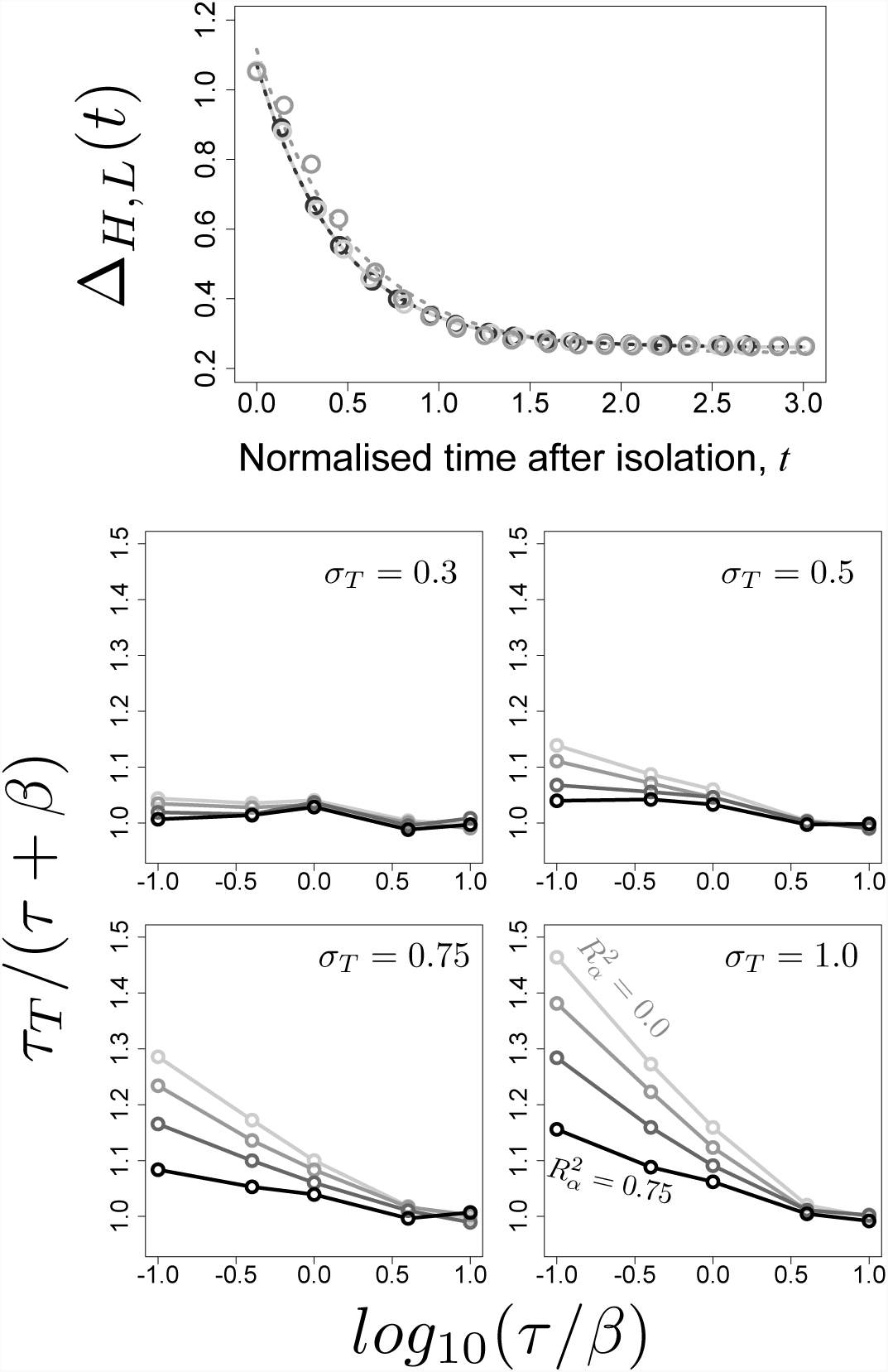
The function Δ_*H,L*_(*t*) decays with approximately exponential dynamics. a) Simulations of the isolation of cells were done, for various values of *τ* and *β*, with 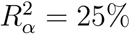. Shown are simulation results (symbols), along with the results of fitting the model of exponential decay Δ(*t*) = *a*+*b* exp (*-t/τ*_*T*_) to the simulation data (dashed lines), where *a* and *b* are constants. Time is normalized in each case by the instant *t*^*\**^ such that Δ_*H,L*_(*t*^*\**^) has decayed by 90%. The light to dark gray tones correspond to the values of the ratio *τ/β* = 0.1, 1.0, 10.0 respectively, with *β* = 50. b-e) Comparison between *τ* + *β* and the value estimated for *τ*_*T*_. Simulated data (Δ_*H,L*_(*t*)) were fitted under the same setup as in (a) and the resulting values of *τ*_*T*_ plotted as a function of the value of *τ* + *β*. Each graph corresponds to simulations using the indicated value of *σ*_*t*_ with different values of 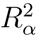 (0.0, 0.25, 0.50 and 0.75) depicted in different gray tones (the darker the tone the higher the value of 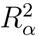).

It follows that the effective characteristic time *τ*_*T*_ is undefined in the case of 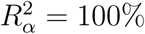, since Δ_*H,L*_(*t*) does not change as a function of time after isolation. Since *τ*_*T*_ is a measure of the time needed for the initial difference Δ_*H,L*_(0) to reach the asymptotic value lim_*t→∞*_ *{*Δ_*H,L*_(*t*)}, it provides a formal characterisation of the timescale of the variation.

An exhaustive simulation study (figure 4, b-e) led to the conclusion that *τ*_*T*_ can be approximated, with a typical bias of at most 5–10% of the true value, as:

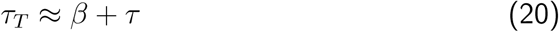

Therefore, the auto-correlation time of the stochastic rate of protein production (*τ*) and the mean lifetime of the protein (*β*) determine the timescale of the variation in expression levels (*τ*_*T*_).

The relative contribution of the stable component 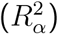 and the effective characteristic of the variation (*τ*_*T*_) can be visualized in a single plot, derived from equation 19. As shown in figure 5, 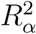 corresponds to the asymptotic value of Ω_*H,L*_(*t*), while *τ*_*T*_ corresponds to the instant of time that satisfies:

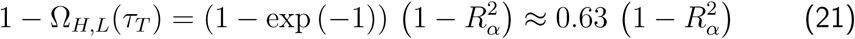

**Figure 5:**
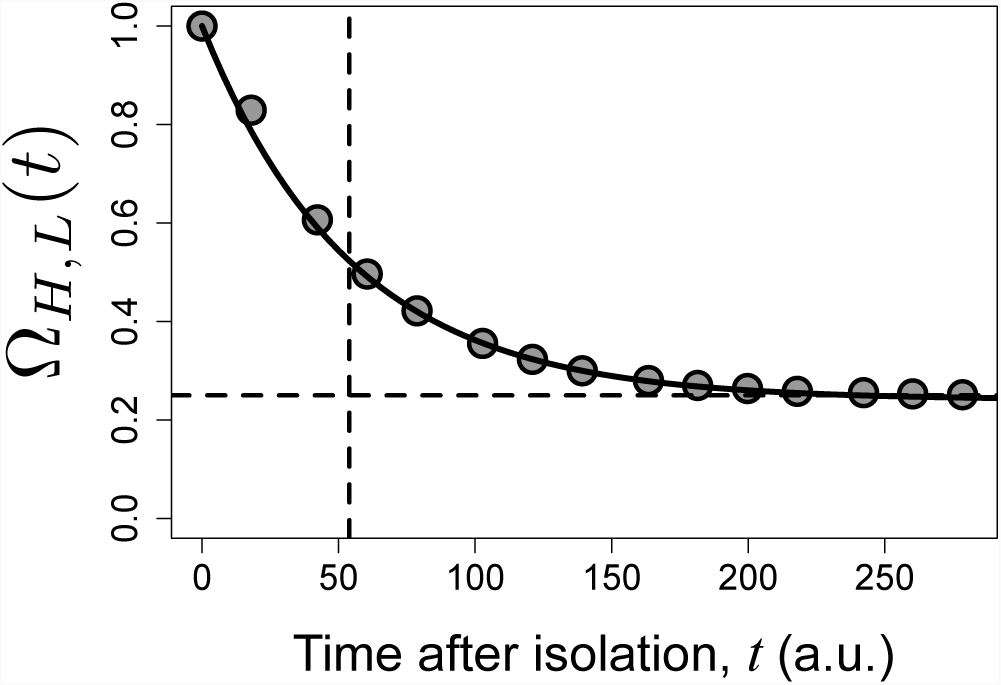
Illustration of function Ω_*H,L*_(*t*). Shown are simulation results (symbols), with 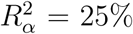, *τ* = 50 and *β* = 5, which were fitted to the expression for Ω_*H,L*_(*t*) in equation 19 (continuous line). The horizontal and vertical dashed lines indicate respectively the true value of 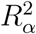 and the value of *τ*_*T*_, as given by equation 21.

Since equation 19 features an exponential decay, it follows that the plateau is reached in practice after an amount of time approximately equal to 5 *τ*_*T*_… Furthermore, the inequality in equation 17 becomes:

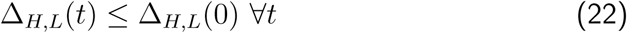

since function Δ_*H,L*_(*t*) is monotonically decreasing with time.

Although this section has focused on the case in which high and low expressors are used, all the properties derived also hold for any two isolated cohorts *A* and *B*. The only requirement is that the condition Δ_*A,B*_(0) *≠* 0 is satisfied.

## 5 Quantification of the components shaping the variation in T-cell receptor expression levels

The theoretical framework developed in the previous section is used here in the analysis of the variation in the expression of the TCR in mouse CD4+ T lymphocytes. The TCR is a heterodimeric membrane receptor that elicits signal transduction upon interaction with MHC-peptide complexes on the membrane of antigen-presenting cells [33]. In wild-type animals, the T cell populations are genetically heterogeneous at the level of the TCR. The genetic diversity of the receptor is brought about by the somatic recombination at the loci encoding the receptor chains in thymocytes (reviewed in [34]). In contrast, genetically manipulated mouse strains are available in which all the T cells express the same TCR (for example, [27]). In these mouse strains, the somatic recombination is ablated (*Rag2*^−/−^ background) and a single functional TCR is expressed in all cells driven by transgenes encoding the two chains of the TCR.

To quantify the origin and timescale of the variation in the context of the TCR, we used a polyclonal population from a wild-type imbred strain and a surrogate monoclonal population from the Marilyn TCR-transgenic strain [27]. In this setup, we are interested in comparing the values of 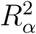 and *τ*_*T*_ estimated for the polyclonal and the Marilyn monoclonal populations. These two populations show comparable mean expression values (see figure 6, top graphs) but the expression is more variable in the polyclonal population than in the monoclonal one [35], presumably reflecting the genetic diversity [36].

**Figure 6:**
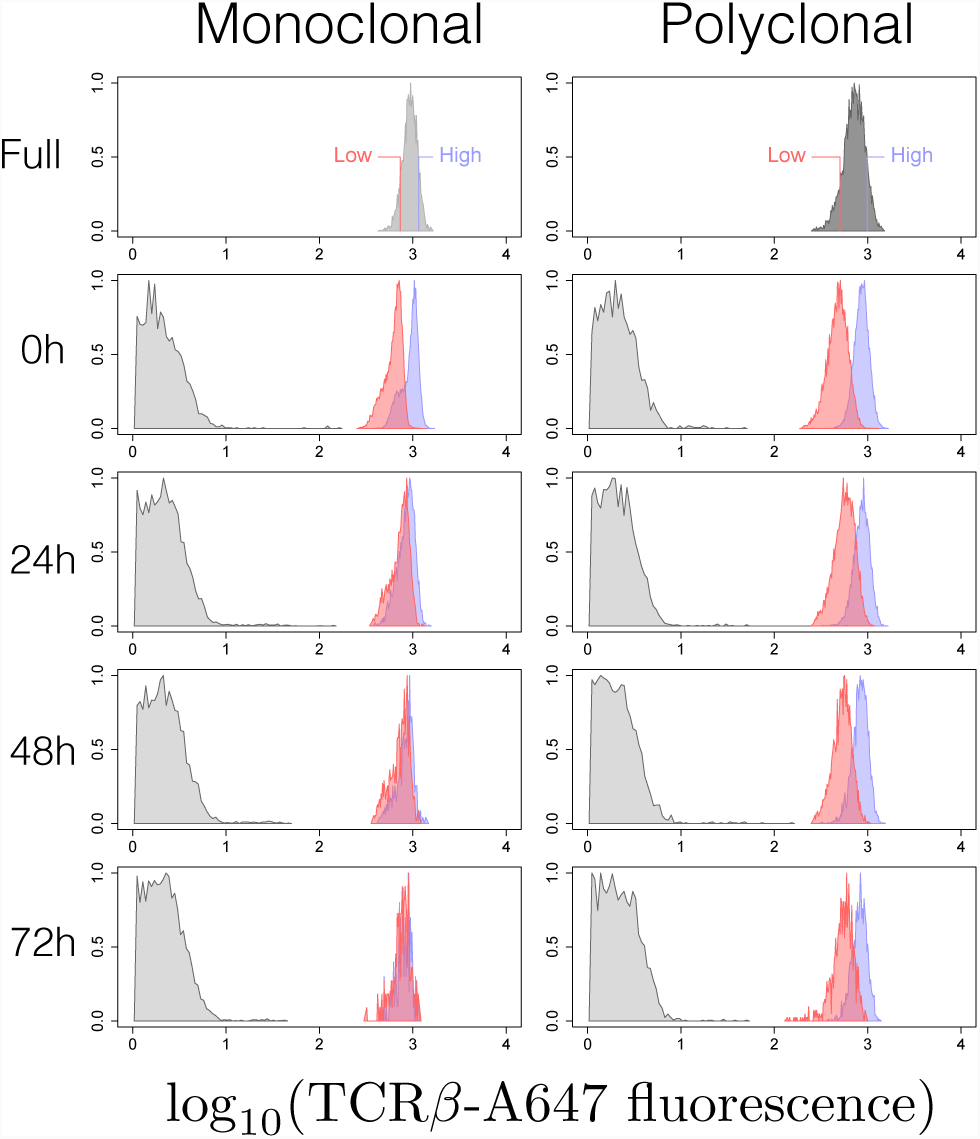
Dynamics of TCR expression in high and low expressor cohorts sorted from monoclonal (Marilyn; left) and polyclonal (wildtype; right) populations. The graphs are the histograms of frequency of log-transformed TCR fluorescence in the high (blue) and low (red) expressors measured by flow cytometry at the indicated times after sorting. Unstained population is also shown.

In the model developed in section 2.2, the stable component arises from different mean protein production rates. In polyclonal populations, the stable variation in average TCR production may be caused by the differential regulation of expression of the receptor sub-units, depending on the specificity of the particular TCR, or by the differential ability of the specific sub-units to pair and be expressed [37]. In either case, genetic heterogeneity would ultimately explain some of the variation observed at the level of a polyclonal population. If so, this would imply in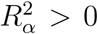 for a polyclonal population. By analysing a TCR-transgenic population, we address whether genetic variation is the only factor explaining the stable component. In the affirmative case, one would obtain 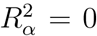 for a TCR-transgenic population. If one obtains 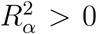, non-genetic mechanisms must be evoked.

We adopted an experimental design in which high and low expressors, defined to contain 10% of the mass of the starting population distribution, were sorted (figure 6, top) and then maintained *in vitro* without any stimulation. As described before [38], there was no cell division under these conditions, and cells slowly died off such that after 3 to 4 days no live cells were left. Since in the Marilyn transgenic strain, all T cells have a naive phenotype [27], we restricted the analysis of the wildtype polyclonal populations to those cells that express high levels of the CD45RB marker, indicative of a naive phenotype [39]. By restricting the analysis to naive cells, the distribution of cell size as measured by Foreward Scatter was similar in the high and low expressor cohorts when sorted from the monoclonal Marilyn population as well as from polyclonal population (supplemental section F).

The dynamics of the frequency distribution of the TCR expression levels in cohorts of high and low expressors sorted from polyclonal and monoclonal animals and subsequently cultured *in vitro* for up to 72h is illustrated in figure 6 for a representative experiment. The distributions of the TCR expression levels in the high and low expressors sorted from wildtype polyclonal population remain clearly different. In contrast the high and low expressors from the monoclonal Marilyn TCR-transgenic population become very similar as a function of time after sorting. In fact, this particular data set portrays an extreme case of convergence of the histograms of high and low expressors and it was chosen to better illustrate the difference between the monoclonal and polyclonal populations. In the two other independent sets, the histograms of Marilyn high and low cohorts did not converge (see figure 7 for the quantification including all experiments).

**Figure 7:**
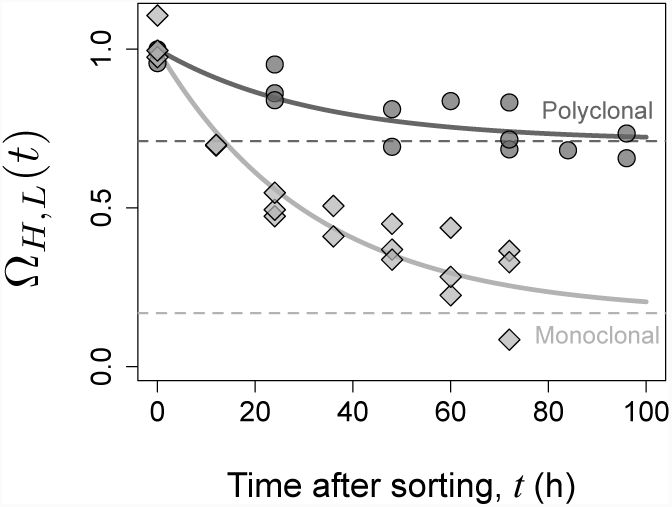
Dynamics of Ω_*H,L*_(*t*) in sorted cohorts. The symbols are the point estimates of Ω_*H,L*_(*t*) = Δ_*H,L*_(*t*)/Δ_*H,L*_(0) at different times after sorting for polyclonal (circles) and monoclonal (diamonds) cell population data sets. The curves represent the best fit of the function Ω_*H,L*_(*t*) as defined by Modelling Scenario 2 to the ensemble of the populations data sets. The horizontal dashed lines indicate the estimates of the asymptotic 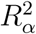.

The values of Ω_*H,L*_(*t*) in three experimental data sets are shown in figure 7. The normalisation (equation 18) masks the fact that the value of Δ_*H,L*_(0) was conspicuously greater for the polyclonal population, in accordance with the observations [35], that the variance 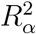 is larger in polyclonal populations than in TCR transgenic populations (figure 8).

**Figure 8:**
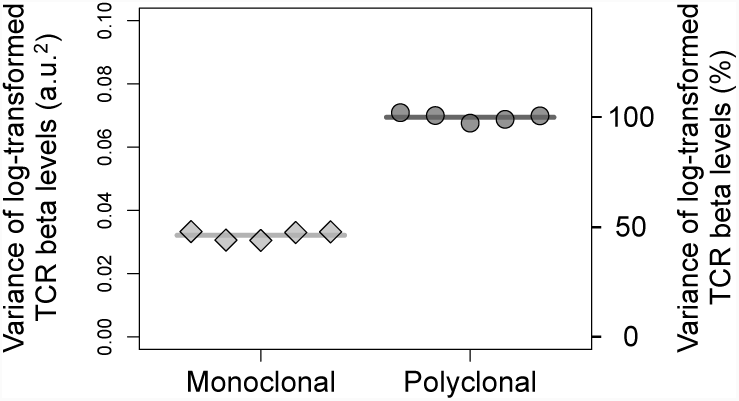
Variance of the log-transformed TCR expression levels in monoclonal (Marilyn; diamonds) and polyclonal (wildtype; circles) CD4+ T lymphocyte populations. The points are estimates of the variance in independent samples and the lines are the average value of these variances.

In order to estimate 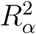 and *τ*_*T*_ by fitting the model to the experimental data equation 19 must be refined as follows:

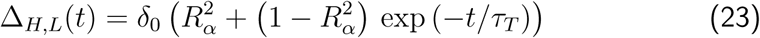

where *δ*_0_ represents an estimate, obtained via fitting, of the “true” initial value Δ_*H,L*_(0). Equation 23 has the important property of preserving the statistical independence between data points used as input for the fitting, a key requirement for proper statistical analyses.

The analysis was based on fitting the three-parameter exponential model (equation 23) to the ensemble of the data, composed of the multiple experiments done for each biological population. The different modelling scenarios being tested are defined by specifying each of the three parameters, 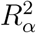, *τ*_*T*_ and *δ*_0_, for each biological population as being shared or not between the polyclonal and monoclonal populations. Small variations in defining the percentages for sorting high and low expressors in different experiments are expected to sporadically affect the value of *δ*_0_ and therefore this parameter was always fitted separately for each experiment. Therefore, the modelling scenarios are obtained by specifying how parameters 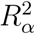 and *τ*_*T*_ are shared between the biological populations. The complete description of the modelling scenarios considered is presented in table 2 (column 2). Modelling Scenario 1 represents the null model, according to which the polyclonal and monoclonal populations are described by the same values of 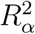 and *τ*_*T*_. This is the scenario with the smallest number of parameters considered. Scenario 2 represents the plausible situation in which these two populations may be described by different values of 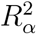, but equal *τ*_*T*_, while in Scenario 3 parameter *τ*_*T*_ is also allowed to be different in the two populations. Finally, Scenario 4 represents a lower bound in terms of the error in the fitting, where data from each experiment is fitted independently, and has the largest number of parameters. The Akaike Information Criterion (AIC) [40] is used to compare the different modelling scenarios in their capacity to fit to the ensemble of the data. The AIC has a solid foundation on information theory [40], representing a compromise between the error in fitting the data and the number of parameters in the model. The results are presented in terms of the difference ΔAIC_*c*_ between the AIC for each Scenario and that of Scenario 1. In comparing different modelling scenarios, the one with the smallest value of the AIC (and therefore, the smallest value of ΔAIC_*c*_) provides the best and most parsimonious description of the data.

**Table 2:** Overview of the Modelling Scenarios tested, with a description of how parameters 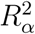 and *τ*_*T*_ were set in the two biological populations, and the resulting number of parameters that are fitted. As discussed in the text, parameter *δ*_0_ were fitted separately for each experiment

| Scenario | Description | # Parameters Fitted |
| --- | --- | --- |
| 1 | $R_\alpha^2$ and $\tau_T$ have the same values in the two biological populations | 8 |
| 2 | $R_\alpha^2$ may have different values for each biological population, but $\tau_T$ has the same value | 9 |
| 3 | Both $R_\alpha^2$ and $\tau_T$ may have different values for each biological population | 10 |
| 4 | Each experiment was fitted independently | 18 |

The results of the model fitting, including the sum of squared residuals, point estimates of the parameters, and the value of ΔAIC_*c*_, are shown for each scenario in table 3. It follows that Scenarios 2 and 3 have the lowest values of ΔAIC_*c*_, with Scenario 3 having a slightly higher value. Since the latter has one extra parameter, this suggests that a scenario where only 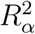 is allowed to be different constitutes the most parsimonious explanation for the data. Hence, altogether Scenario 2 is favoured, according to which the two populations differ only in 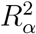. In this case, we obtain an effective timescale of 32 hours, and values of 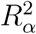 of 71% for the polyclonal and 17% for the monoclonal population, with 95% confidence intervals of [57%, 79%] and [0%, 31%], respectively. Finally, the function Ω_*H,L*_(*t*) resulting from scenario 2 is shown in figure 7, highlighting the values of 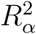 estimated for each population.

**Table 3:** Estimates for the parameters of the populations obtained by fitting the data on Δ_*H,L*_(*t*), based on the different modelling scenarios under consideration. The results are presented in terms of ΔAIC_*c*_, the difference between the value of the AIC (corrected for small sample size; see methods) of each scenario and scenario 1. Modelling scenarios with lower values of ΔAIC_*c*_ provide a more parsimonious explanation for the data

| Fitting $\Delta_{H,L}(t)$ | | | | | | | |
| --- | --- | --- | --- | --- | --- | --- | --- |
| Scenario | SS Residuals | Population | Exp. # | $\delta_0$ | $R^2_\alpha$ (%) | $\tau_T$ (h) | $\Delta AIC_c$ |
| 1 | 0.103 | Polyclonal | 1 | 0.55 | 61 | 32 | 0.0 |
|  |  |  | 2 | 0.62 |  |  |  |
|  |  |  | 3 | 0.63 |  |  |  |
|  |  | Marilyn | 1 | 0.24 |  |  |  |
|  |  |  | 2 | 0.29 |  |  |  |
|  |  |  | 3 | 0.26 |  |  |  |
| 2 | 0.028 | Polyclonal | 1 | 0.52 | 71 | 32 | -40.5 |
|  |  |  | 2 | 0.58 |  |  |  |
|  |  |  | 3 | 0.59 |  |  |  |
|  |  | Marilyn | 1 | 0.37 | 17 |  |  |
|  |  |  | 2 | 0.40 |  |  |  |
|  |  |  | 3 | 0.37 |  |  |  |
| 3 | 0.025 | Polyclonal | 1 | 0.51 | 59 | 75 | -39.6 |
|  |  |  | 2 | 0.57 |  |  |  |
|  |  |  | 3 | 0.58 |  |  |  |
|  |  | Marilyn | 1 | 0.37 | 24 | 24 |  |
|  |  |  | 2 | 0.41 |  |  |  |
|  |  |  | 3 | 0.38 |  |  |  |
| 4 | 0.015 | Polyclonal | 1 | 0.50 | 22 | 199 | 2.2 |
|  |  |  | 2 | 0.57 | 67 | 46 |  |
|  |  |  | 3 | 0.57 | 0 | 249 |  |
|  |  | Marilyn | 1 | 0.40 | 4 | 32 |  |
|  |  |  | 2 | 0.39 | 40 | 16 |  |
|  |  |  | 3 | 0.37 | 25 | 27 |  |

## 6 Discussion

In this article, we developed a new approach to analyse the variation in protein expression levels in a cell population, which enables measuring the characteristic dynamics of the fluctuations in cellular levels and estimating the magnitude of stable and unstable contributions to the variation across cells. The crux of the proposed analysis is based on the realisation that the difference between the means of log-transformed expression levels in two selected cohorts isolated from a population of interest converges with aproximmate exponential dynamics to an asymptotic value. By normalising this asymptotic value by the difference in cohorts’ means immediately after their isolation one obtains an unbiased estimation of the proportion of population variance that is explained by the stable component 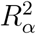, while the mean convergence time *τ*_*T*_ measures the timescale of unstable component dynamics. This is the key insight of the analysis. This insight stems from perceiving any cell population as a mixture of many independent subpopulations, each with a characteristic mean expression level, that is fixed yet distributed among the subpopulations. Under these assumptions, the population variance is equated to the sum of the variance of the subpopulations means, which embodies the stable component of variation, and the variance of the expression level within the subpopulations, which represents the unstable component.

At first sight, the stable and unstable components of expression variation as formulated here are analogous to what Huang [10] referred to as population and temporal noise, respectively. However, this analogy is not straightforward. Huang’s definition of population noise precludes, by construction, any underlying genetic and stable epigenetic mechanisms. In contrast, the stable component, as defined here, is a statement about the dynamical aspects of variation and is silent about mechanism. We believe that the terms stable and unstable components are not only intuitive but convey a more precise description of variation in terms of its temporal dynamics. The mechanistic basis of these components remains a matter for further analysis. Putative mechanisms underlying the stable component include genetic variation and non-volatile epigenetic traits [41, 10]. In turn, the unstable component may be explained by noise in gene expression [9, 10] as wells as transient metastable epigenetic variants [41, 10]. Stable gene expression variants, which would be part of stable component of variation, are expected to be pervasive, since differentiation stages, cell lineages and cell types are hallmarks of multicellular organisms [25, 26]. In spite of this expectation, many quantitative approaches to expression variation in cells in the past have focused on noise in gene expression [4, 19, 7, 21, 23, 24].

Measuring the extent to which selected cohorts of cells can restore the full complexity of population from which they were sorted is an intuitive approach to analyse the heterogeneity of a population. This basic intuition motivated the experimental design used in several reports [16, 42, 14, 17, 18], in which the stable and unstable components of variation were evoked and utilised in an informal way. The capacity of cohorts to restore totally or partially the distribution of the original population has often been interpreted and discussed qualitatively, based on the visual inspection of raw flow cytometry histograms or of summary data time series. The present report advanced beyond such ‘half-full / half-empty glass” interpretations of data by contributing a rigorous quantitative method to analyse these kind of sorted cohort experiments based on estimating two parameters, 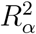 and *τ*_*T*_, that capture the essence of heterogeneity and dynamics of the expression variation.

The analysis method is grounded on a stochastic modelling framework. The protein expression levels in a single cell are described as very simple stochastic processes, based on [29], in which the instantaneous protein production rate (captured by variable *z*_*t*_) fluctuates generating a stationary log-normal distribution of expression levels in each subpopulation. Protein expression has been modelled by others considering transcriptional burst dynamics that can be shown to generate discrete numbers of transcripts following a negative binomial distribution. Although transcript copy number distributions are generally assumed to be described by a negative binomial distribution at any range of expression levels, they are well-approximated by a log-normal distribution at high copy numbers per cell [43]. Therefore the log-normal approximation underlying the dynamics of *z*_*t*_ is justified by the observation that the transcripts encoding the TCR *α* and *β* chains are among the most abundant in the cell [44]. This model of the single cell expression dynamics was used to simulate a population formalised as a large mixture of independent subpopulations. By applying the equation for partitioning the variance to this mixture model, the analysis based on log-transformed values emerged as the best approach, since in this case the contributions due to the stable and unstable components are additive, greatly simplifying inference. This is particularly relevant for flow cytometry data, which is typically analysed in a logarithmic scale. It is interesting to note that [18] also relied on log-transformed values for quantification, based on the analysis of time-series of expression levels in individual cells.

An important result here is that the rigorous unbiased estimation of 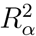 can be done based on a time series of normalised measurements of the difference between the means of log-transformed expression levels in isolated cohorts. The normalisation of these measurements by the value immediately after sorting is a critical part of the inference procedure. A similar normalisation by the initial value was used by Singh [31] to analyse the temporal evolution of the squared coefficient of variation of a single population, under a model that assumed that the observed variation was completely due to noise in gene expression. Using this type of analysis in settings of transcription inhibition, these authors [31] assessed whether noise in mRNA production and degradation or promoter activity fluctuations contribute to noise in protein expression. The normalisation of the differences by their initial value (*t* = 0) in the present work formalises the definition of how much of the initial difference, introduced by the process of sorting by design, remains at later times (function Ω(*t*)). Hence, a key requirement is that the isolated cohorts being compared have different means just after sorting. This strongly argues to using high and low expressors as the basis for quantification, in order to maximise the measurement resolution. In practice, one has to manage a tradeoff between how extreme are the expression levels (to increase resolution and dynamic range of the readout) and how many cells are contained in the cohorts (sample size). The estimation based on the mean expression levels offers great advantages in terms of the range of applications. It is often argued that the standard experimental techniques that measure bulk expression are obsolete in the context of the studies of gene expression noise, because their population-averaged readouts mask cell heterogeneity (see, for example, [10, 45]). The present analysis framework enables to use these techniques, as one may combine the isolation of cells (the only step requiring the analysis at the single-cell level), with population-averaged readouts to quantify 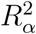 and *τ*_*T*_. The function Δ(*t*), which is at the core of the estimation process, can be approximated as the logarithm of the fold-ratio between the raw mean values of the two populations. In theory, by measuring 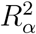 and the total variance of the full population 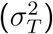, one could estimate the actual values of 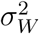 and 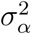 in equation 13. This would allow one to compare the values of 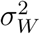 and 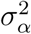 in different biological populations Furthermore, we also showed that the variances of the isolated cohorts can be further informative, allowing the estimation of the ratio between the absolute values of the contribution of the stable component in the isolated cohort and in the starting population. However, the estimate obtained in this way is biased, under-estimating the true value by up to 20%. Consequently, if an estimate of this ratio is needed, we suggest a simulation-based approach.

Empowered by the quantitative framework, we analysed the variation in the expression levels of the T-cell receptor (TCR) in mouse CD4^+^ T cells. T cells are an interesting case study. It has been described that some molecules in these cells have a component of stability in their expression levels, such as CD5 [46, 47, 48]. This has also been suggested for CD127 [49], a sub-unit of the IL-7 receptor. Also studies centred on chromatin regulation of expression of cytokines in T cells [13, 15], found that the expression of IL-4 and IL-10 is unstable at the level of single cells. With the increasing availability of single cell genomics, proteomics and metabolomics techniques there is accumulating evidence that cell populations that hitherto were perceived as homogeneous are in fact complex mixture of cell types and cell states, that may be reversible and transient, raising the issue of stability and dynamics. From a practical perspective, different mouse models are available with different genetic diversity in the TCR loci, which gives a handle to tease apart genetic and non-genetic components of variation. Hence, in our analysis of TCR expression levels we studied a genetically heterogeneous polyclonal population, and also a particular isogenic population, Marilyn TCR transgenic [27] in a *Rag2*-deficient background. These two populations display distinct variances of the TCR levels that are, not surprisingly, positively associated with the genetic TCR diversity. We asked whether these two populations could be described by equal or different values of 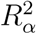 and *τ*_*T*_ and found that the overall most parsimonious explanation for the data was a model where only one of these parameters is allowed to be different. The model with different *τ*_*T*_ and the same value of 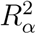 performed marginally better based on the AIC. However, the point estimate of the characteristic time of the polyclonal population in this scenario was about 20 days, requiring an extrapolation beyond the experimental observation time of 96 hours and involving an unreasonable large uncertainty. Although such long time scales have been described for the restoration of a bimodal population distribution from selected unimodal cohorts (e.g. over 30 days in [18, 42]), the scenario of very distinct *τ*_*T*_ values for wildtype and transgenic populations is biologically unsound. This scenario requires that the expression of transgenic TCR would differ from natural TCR expression in terms of the protein turnover rate *β* as well as noise in gene expression, such that the same 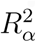 could be present in populations with markedly different variances (figure 8). Therefore, based on these statistical and biological considerations we rejected this scenario. We concluded in favour of the scenario in which the TCR expression fluctuates with a characteristic time of 32 hours in the both populations, which differ in the values of 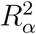, the polyclonal having 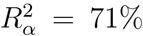and monoclonal Marilyn having 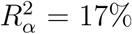. The relatively small yet not negligible value of 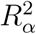 obtained for the latter population may be particular and not necessarily generalisable to other TCR-transgenic populations. It is worth mentioning that the analysis of another such TCR transgenic population led to a higher 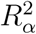 value [50], suggesting that transgenic populations, which are known to display different variance of the TCR levels (e.g. [35]), may also differ in the extent of the stable component of variance.

The capacity to analyse the variation in TCR expression validates experimentally the theoretically-designed methodology. We could quantify the two key parameters in the two cell populations, implying that the methodology has enough power to resolve the stable and unstable components of variance even when the distribution of interest is remarkably unimodal and narrow.

Beyond this key methodological result what do the actual estimates of 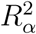 and *τ*_*T*_ tell about phenotypic variation in TCR expression?

It is important to highlight from the outset that the stable proportion of variance and the timescale may be constrained by the experimental setup on which we intentionally relied. High and low expressors were maintained *in vitro* for as long as possible in the absence of any stimulation in conditions that precluded cell division. Using this setup, we focused on cell-intrinsic components only, avoiding complications arising from cell division. As a consequence of this choice, the present data do not exclude the possibility that signals arising from the intermittent stimulus from the antigen-presenting cells in the *in vivo* environment may change the values of both 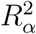 or *τ*_*T*_ for the populations tested. Also, cell division is expected to decrease the timescale of the fluctuations in a twofold manner. First, protein dilution into the daughter cells may effectively reduce the value of *β*, even if yeast studies indicate that protein levels are remarkably constant if corrected for cellular volume [51]. Second, cell division may affect the stability of epigenetic modifications facilitating the transitions between chromatin states or bistable transcriptional switches that affect quantitatively TCR expression in this way reducing the effective *τ*_*T*_. A similar point was made in a study [52] of induced pluripotent stem cells. These cells maintained a memory of transcriptional and epigenetic signatures indicative of the cell of origin that vanished with sequential passages. Hanna et al. [53] reported a similar impact of cell division itself. Furthermore, cell division and generation time variability may introduce cell-extrinsic deformations of the expression distribution by differential selection of lineages (see [54] for a theoretical analysis). These potential peculiarities of the experimental design notwithstanding, the estimates of 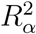 and *τ*_*T*_ are to our knowledge the first reported values and therefore interpreting the meaning of these values requires indirect comparison with other estimates.

The characteristic time of the variation in protein expression represents a transient memory of expression levels [7]. Various studies have quantified the dynamics of fluctuations in expression levels of various molecules, reporting characteristic times that range from hours [7, 12] to days and weeks [16, 14, 17, 18, 42]. In studies quantifying the dynamics of the percentage of T cells inducing the expression of cytokines [13, 15], the effective timescale was estimated to be about 70 hours for the cytokines IL-10 [13] and IL-4 [15], under conditions in which the cells divide. In terms of molecular mechanism, this has been linked to slow dynamics of chromatin remodeling [13, 15]. The longer time scales were systematically obtained in scenarios with cell division and that involved the restoration of a multimodal distribution of expression levels from biased cohorts. The dynamics of multimodal distributions, in which cells switch between overtly distinct subpopulations may correspond to transitions between cellular states. For the unimodal TCR expression, we estimated an effective timescale of 32 hours, in the absence of cell division. This timescale is shorter than that necessary to restore a full multimodal distribution from extremely biased cohorts [18, 42]. The TCR protein complex is arguably one of the most complex receptors in terms of its composition, trafficking and regulation. In quiescent cells, such as the naive cells analysed here, it is continuously recycled between the plasma membrane and intracellular membranes with a fast rate of less than an hour. The TCR in the ensemble of these two pools has a slow turnover rate. The treatment with protein synthesis inhibitor up to 12 hours led only to modest changes in expression levels [55], suggesting that *β* might be greater than 12 hours. However, this estimate is potentially problematic, since this treatment may alter the regulation of the TCR complex levels, as it may up-regulate the expression of the mRNA encoding its *ζ* chain sub-unit [56]. Sousa and Carneiro [57] estimated the baseline TCR turnover in an human T cell line by fitting the dynamics of the mean upon short-term stimulation, and found a value for *β* of 15 hours. Both values [55, 57] are compatible with the effective timescale estimated here, which lumps protein stability and the auto-correlation time of the rate of protein production, and suggest that *β* is of the same order of magnitude as *τ*_*T*_ in the case of the TCR.

The different components that may underly the stable variation in TCR expression levels are systematically addressed in figure 9. The mean TCR level has been shown to be distributed among the V_*β*_-family subsets in CD4 as well as CD8 human T cell populations [36], under conditions in which there was a strong correlation with cell size. Despite the fact that the high and low expressors cohorts in our experiments had virtually the same size distribution as assessed by the respective forward scatter signal, it is likely that part of the stable variance in TCR expression is in fact due to the genetic diversity at the TCR locus. The question is whether genetic diversity explains all stable variation. The estimate of a positive value for 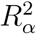 in the nominally monoclonal TCR transgenic population suggests that non-genetic variation may contribute to stable differences in expression levels of the TCR among cells. This might be a particular feature of TCR-transgenic populations, as their relationship to the actual clones in a polyclonal T cell population is not trivial. The specific mechanism that would mediate such non-genetic variation is unclear at present. We speculate that the stable component in this system may arise from a myriad chromatin modifications in the form of “molecular switches”, which would affect, directly or indirectly, the expression of the TCR. This speculation is inspired on theoretical studies [58, 13], which have predicted these marks to be stable once fully established, but also potentially variable among cells. In terms of the full range of modifications affecting expression of the TCR in *cis* and in *trans*, some could be present, while others could be absent in each individual cell in a stochastic yet stable pattern of modifications [58, 13]. In those cells in which the balance of modifications happens to be tilted towards those inducing expression, levels of the TCR would be higher than average, while in cells with lower TCR levels this balance would be shifted in the direction of those leading to decreased expression. Similar considerations could be made to any epigenetic variants in any of the vast number of transcription factors and regulatory proteins that control the TCR complex expression. Finally, in this enumeration of the causes of stable variation in TCR levels, it is worth noting that despite we have been referring to the population of T cells in Marilyn transgenic mice as “monoclonal” throughout this article, the cells are not a T cell clone derived from a mature T cell. Instead, they are continuously differentiating in the thymus and being exported to circulation. One cannot rule out that these T cells or their bone marrow and thymic precursors underwent sporadic somatic mutations in any of those genes affecting TCR complex expression. The existence of genetic mosaics in somatic tissues has been well documented (reviewed in [59]) following the advent of single cell sequencing, and thus, one must envisage the possibility that part of the stable variation observed in transgenic TCR expression is due to *bone fide* genetic variation. Simple back of the envelop calculations suggest that epigenetic variants and/or mutational mosaicism outside the TCR locus may represent more than 1/6 of the stable variance in the wild type CD4 T cells (assuming that epigenetics and/or mosaicism explain the same amount variance in both monoclonal and polyclonal populations, that all the stable variance in the former (which corresponds to 17% relative to monoclonal and 8% relative to polyclonal variances) is explained by these two processes and that the stable variance in latter is explained by these two processes and by TCR diversity, we have that in the polyclonal population 8/71 of the stable variance is explained by epigenetics/mosaicism and 63/71 is explained by genetic TCR diversity; see figure 9).

**Figure 9:**
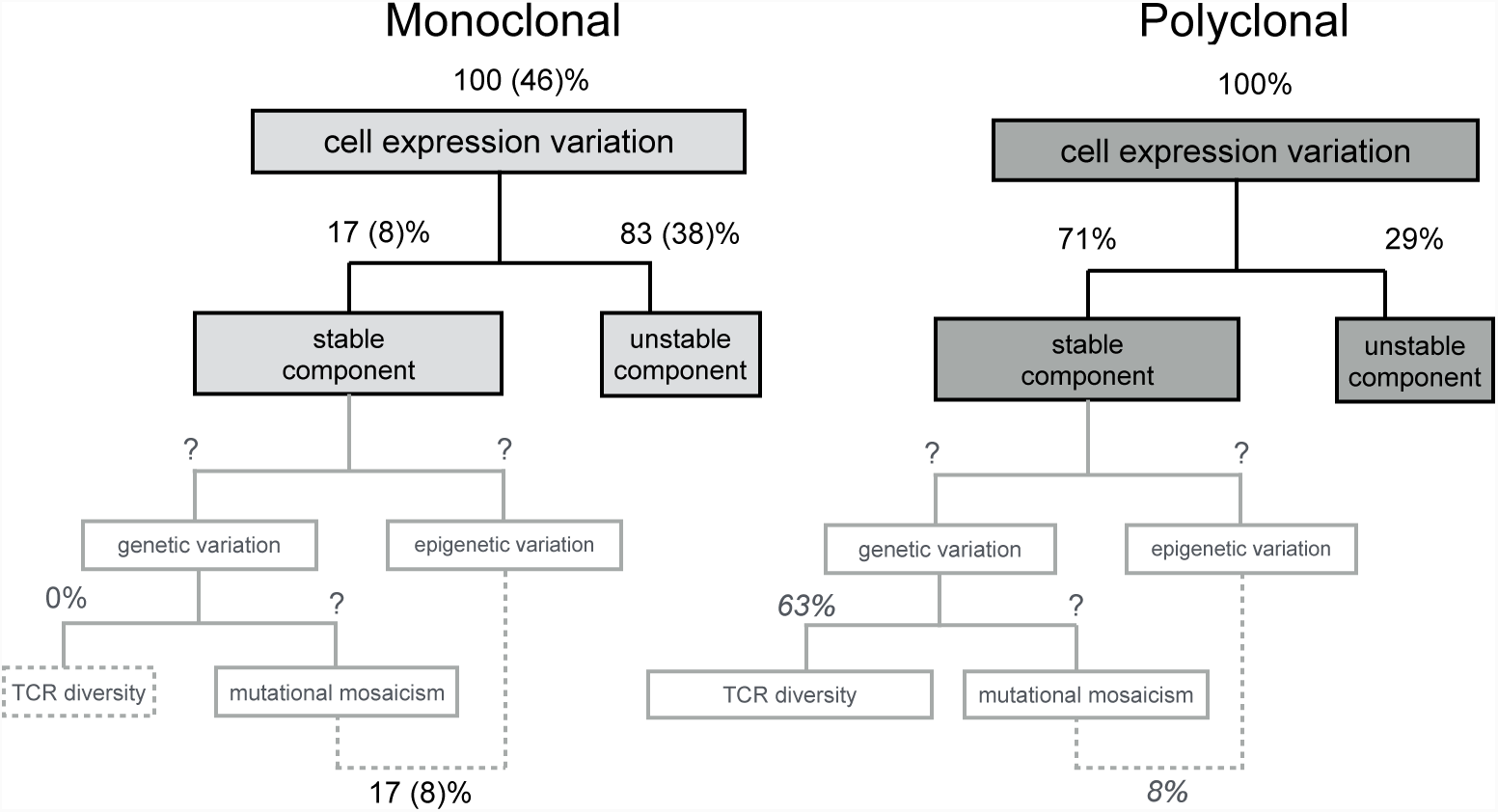
Overview of the components of TCR expression levels variation in naive CD4 T cells from monoclonal Marilyn transgenic and polyclonal wildtype mice. The first partition of the variance in each population corresponds to the stable and unstable components experimentally estimated in this article. The further partitions of the stable component are indicative of the putative genetic and epigenetic causes of the variation. The percentages represent the expected proportions of the variance in log transformed TCR expression levels explained by the indicated components. The values in brackets in the diagram of the monoclonal population were normalised by the variance in the polyclonal population. The values in black are experimental estimations. The values in grey italic are guesses obtained by assuming that the variance explained by somatic chimerism and/or epigenetics is the same in polyclonal and transgenic populations.

This quantitative framework makes a connection between systems biology, in particular works focused on noise in gene expression [9], and quantitative genetics [60]. In both domains, approaches for decomposing the variance, or another measure of variation, have been instrumental in studying the properties of different biological systems (see, for example, [41]). In the studies of noise in gene expression, the notion of intrinsic and extrinsic noise put forward by Elowitz and colleagues [4, 19] is based on decomposing the coefficient of variation of expression levels into these two noise sources. Other works have focused on either generalising this distinction or developing new decompositions [21, 61, 24, 23]. In fact, these approaches may be combined with the framework developed here, to further partition the unstable component, for example, into intrinsic and extrinsic noise. Likewise, in connection with quantitative genetics, parameter 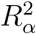 can be interpreted as the “heritability” in expression levels of a population. This arises from an analogy with the decomposition of phenotypic variation into a contribution from additive genetic variation and another due to environment (see, for example, [28, 60]), neglecting non-additive genetic variation. At present, this constitutes a mere analogy, since 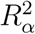 is defined even in the absence of cell division.

Hitherto, the studies on the phenotypic variation in gene expression levels at the individual cell level have relied experimentally on timelapse imaging of single cells or on population snapshots using single-cell resolution techniques such as flow cytometry, qPCR, RNAseq or Cytof. The quantitative framework and methodology proposed here, relying on estimates of the mean of the expression levels in apropriately selected cohorts, enables studying the sources and dynamics of the variation in cellular expression levels by conjugating a single step of sorting with the full gamut of transcriptomic, proteomic, and metabolomic technologies available to measure bulk expression. This opens new prospects for studying quantitative traits and responses in heterogeneous cell populations. In particular, by providing a rigorous approach to quantify the relative contribution of the stable component, this work allows the determination of the extent this component contributes to variation in different biological and experimental systems [16, 14, 17]. Furthermore, the model for expression levels considered here can be further extended to incorporate, for example, more elaborated formulations, such as those with positive and feedback loops, as in the case of gene regulatory networks regulating cell differentiation (for example, [62]). Finally, the sophistication of DNA recording-based methods to account for past fluctuations in transcript levels of a cell or its lineage will generate massive sets of single-cell transcriptomics data points suitable for decomposition into stable and unstable components [63]. In summary, we have put forward a solid theoretical framework to dissect the components of variation in expression levels according to their stability and dynamics, which enables the further analysis of how different molecular mechanisms may modulate each component.

## Materials and Methods

### Notation

The function log(.) denotes the natural logarithm, and random variables are represented as bold symbols, as in ***x***. We use 𝔼 [***x***] to denote the expected value of a random variable ***x***, and 𝕍 [***x***] the variance. The notation ***z*** *∼ℒ 𝒩 (µ, σ*) represents a random variable ***z*** following a lognormal distribution with parameters *µ* and *σ*, having therefore the probability density function:

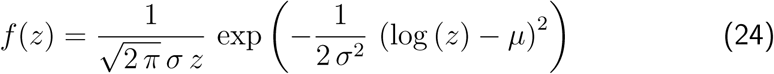

### Numerical simulations

Simulations of the model based on stochastic differential equations were performed using custom software written in C++, based on the GNU Scientific Library (http://www.gnu.org/software/gsl/). For a given value of the parameters *τ* and *β*, the stochastic model (equations 4 and 5) was simulated, using the Brent-Dekker method (GNU Scientific Library) to adjust the value of *σ* so as to obtain the desired value of *σ*_*W*_.

Simulations of cell sorting experiments to isolate appropriate cohorts were done using an initial population with *σ*_*T*_ = 0.3, having 1.2 × 10^6^ cells and 2 × 10^4^ sub-populations, with the number of cells per sub-population following a multinomial distribution. From the starting population, 10% of cells were isolated. As a simple approximation of an experimental setting, each isolated cohort was divided into 3 replicates, and simulated for a given period of time, with snapshots of each replicate being collected at equally spaced times.

### Data analysis, fitting and model selection

Numerical analysis was conducted using MATLAB (Mathworks). The exponential model was fitted to the data by non-linear least squares. To study the relationship between *τ*_*T*_ and parameters *β* and *τ*, simulations were ran for several combinations of values of 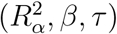. The values of *τ*_*T*_ were estimated by fitting the exponential model. Fitting the ensemble of the experimental data was done by equally weighting each experiment, based on the number of data points per experiment. Values of the Akaike Information Criterion (AIC) were corrected for small sample size, as highlighted in section 2.4 of ref. [40], and include the residual variance as an additional effective parameter being estimated for each model. Confidence intervals (95%) were obtained by bootstrapping each experiment separately, then fitting the ensemble of the data.

### Mice

C57BL6/J and B6.*Rag2*^−/−^ mice were obtained from the Jackson Laboratory. Marilyn mice [27] were kindly provided by Olivier Lantz (Institut Curie, France), and bred with B6.*Rag2*^−/−^ to produce Marilyn.*Rag2*^−/−^. Mice were bred and maintained under specific pathogen free conditions at the animal house of the Instituto Gulbenkian de Ciência, and used for experiments with ages between 8 and 12 weeks. All animal procedures were approved by the ethics committee of the Instituto Gulbenkian de Ciência.

### Antibodies and Flow Cytometry

Flow cytometry was performed using a Beckman-Coulter CyAN ADP. Fc receptors were always blocked prior to staining, by incubation with FcBlock (2.4G2, produced in-house). Cells were stained at 4*°C*, in ice-cold buffer with PBS, 5% fetal bovine serum (PAA), and, except in the case of sorting, with 0.1% sodium azide.

Monoclonal antibodies produced in-house used were: anti-TCR-C*β* (H57-597), anti-CD4 (GK1.5), anti-CD8 (YTS169.4), anti-CD25 (PC61), anti-CD45RB (16A), anti-CD62L (MEL-14), anti-B220 (RA3-6B2), anti-MHC-II (M5/114),anti-Mac1 M1/70), anti-CD3*ϵ* (2C11), anti-CD3*ϵ* (2C11). Commercial antibodies were: anti-CD49b (pan-NK, DX5, BD), anti-CD4 (RM4-5, BD), anti-CD44 (MEL-14, eBioscience), anti-TCR*γδ* (GL3; BD). Biotinylated antibodies were further labeled with PE-Streptavidin (BD).

### Cell sorting and in vitro cultures

Single-cell suspensions were prepared from lymph nodes, and also spleens in the case of Marilyn.*Rag2*^−/−^ animals (due to limited number of cells), by passing cells through a nylon mesh. Cohorts of cells were sorted according to the TCR levels on a FACSAria (BD), using a strategy based on negative selection of CD4^+^ T cells. Briefly, cells were stained for TCR (anti-TCR-C*β*) and lineage markers not expressed by naive CD4^+^ T cells, and then lineagecells falling within the desired TCR gates (illustrated in figure 6 and F) were sorted. A polyclonal naive population was sorted as CD45RB^high^, lineage(CD8, pan-NK, B220, TCR*γδ* and CD25) cells, while Marilyn cells were sorted as CD62L^+^, lineage(B220, CD11c, pan-NK, Mac1, MHC-II). The use of CD62L as an alternative marker of naive cells allows for a more efficient sorting (due to a slow loss in the CD45RB signal throughout the sorting), given the limited number of cells, based on the fact that the vast majority of Marilyn cells retain a naive phenotype [27]. Before each sorting for Marilyn cells, the gating for CD62L^+^ Lineagecells, when analysed in a control sample also labeled for CD4, includes more than 80% of TCR^+^CD4^+^ Marilyn cells. Purities of the sorted populations were assessed by staining aliquots of the sorted populations for CD4 expression, were typically greater than 96%.

After sorting, T cells were cultured in flat-bottom 96-well plates (50 × 10^3^ cells per well), in RPMI (Invitrogen), 10% fetal bovine serum (PAA), 1% Sodium Pyruvate (Gibco), penicillin/streptomycin (Gibco), gentamycin (Sigma), 50*µ*M 2-ME (Gibco), in an incubator at 37*°C*, with 5% CO_2_.

TCR levels were quantified by staining, under optimal, saturating conditions, with anti-TCRC*β* antibody, which binds to the constant region of one of the sub-units of the TCR (see, for example, [64]). Cells maintained in culture were analysed at different time-points by re-staining the TCR, using the same antibody anti-TCRC*β* (clone and fluorochrome) as used for the sorting. In each time-point, 3 replicates (wells) of each sorted population were analysed. In each experiment, an additional population (control) was sorted in parallel, keeping the same gates used for all expressors, but without staining for the TCR, as a control for the impact of this staining. In each time-point, TCR levels of the control population were compared against those of “all expressors”, confirming that the staining for the sorting does not induce massive changes in TCR expression levels.

Data were analysed using FlowJo 8.8.7 (Tree Star Inc.). Cells gated on forward-scatter and side-scatter, live cells (propidium iodide negative) and CD4^+^ cells. For the analysis of TCR levels, cells were further gated on CD62L^+^ cells, to reduce experimental variation in TCR levels. Percentages of CD62Lcells were always lower than 20% in early time-points (up to 48 hours), and similar to those from control cells, arguing against impact of staining for the TCR in order to sort cells. Gated data was exported as text files and analysed in MATLAB (Mathworks) using custom code.

## Acknowledgements

We are grateful to Jocelyne Demengeot and Henrique Teotonio for the support during the development of this work and to Alberto Darszon and Vera Martins for reading an earlier version of this manuscript. We thank Rui Gardner, Telma Lopes and Claudia Bispo for assistance on flow cytometric analysis and cell sorting. This work was supported by a grant from the Fundacao para a Ciencia e Tecnologia (FCT) (PTDC/BIA-BCM/108020/2008). TSG was supported by a fellowship from FCT (SFRH/BD/33572/2008).

## Author contributions

This work was done as a collaboration between all authors. TSG, VMB and JC designed and planned research. TSG developed the theoretical framework, performed simulations, established the experimental setup, performed experiments, analysed data and drafted the manuscript. VMB and JC jointly supervised the work, contributed to both the theoretical and experimental parts, and wrote the manuscript together with TSG. All authors read and approved the final paper.

## Supplementary information

### A Detailed derivation of the mean and variance of the full population

This section presents the detailed derivation of the mean and variance of the full population, given the parameters describing the sub-populations (equations 2 and 3 in the main text).

#### A.1 Mean and variance given the parameters of each subpopulation

We start from the mixture model formulation, where *x* represents protein expression levels:

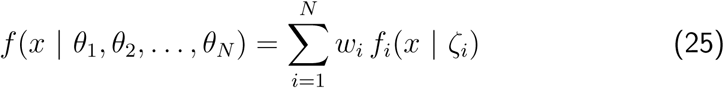

where *θ*_*i*_ = (*w*_*i*_, *ζ*_*i*_) are parameters describing each of the *N* sub-populations. The frequency of the cells in the full population that belong to the *i*-th sub-population is represented by *w*_*i*_ (see equation 1 of the main text), while *ζ*_*i*_ parametrizes the probability density function *f*_*i*_(*x | ζ*_*i*_) of protein expression levels in that subpopulation. For example, in the case that *f*_*i*_ is a normal distribution, *ζ*_*i*_ would be the mean and variance of expression levels in that sub-population.

It follows from equation 25 that the mean of the full population is given by:

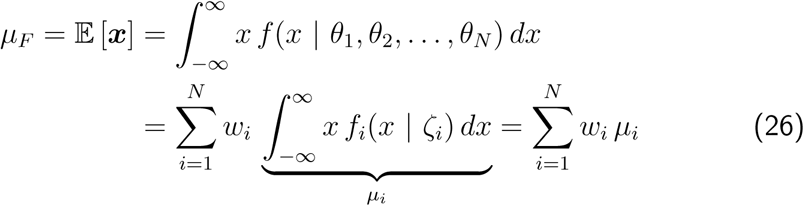

where *µ*_*i*_ is the mean of the *i*-th sub-population. Therefore, the mean of the full population (*µ*_*F*_) is simply the average of the means of the sub-populations, weighted by the frequencies *w*_*i*_. The variance of expression levels, on the other hand, follows from:

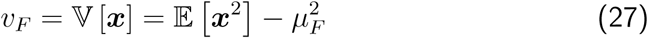

where:

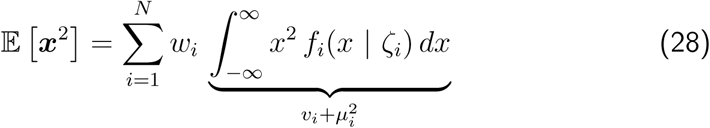

*v*_*i*_ being the variance of each sub-population. Hence, the variance is given by:

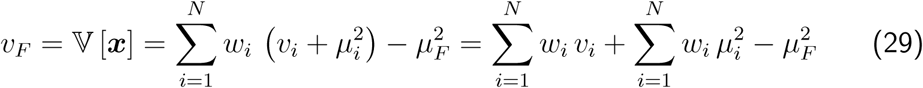

By Jensen’s inequality, it follows that:

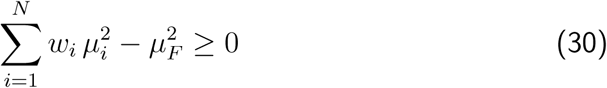

and, therefore, the variance *v*_*F*_ is always non-negative, as expected.

Therefore, for the “full” population, one has the mean and variance given by:

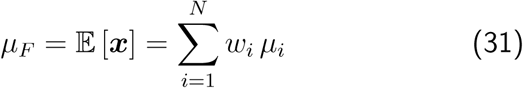

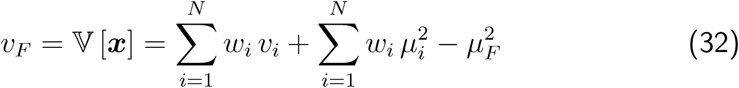

As a remark, these results are independent of the underlying probability density functions *f*_*i*_ describing the expression levels in each sub-population.

#### A.2 Mean and variance in the limit of large number of sub-populations

In the following, we study the asymptotic properties of the equations describing the mean and variance of expression levels in the full population (equations 31 and 32, respectively). In this case, the parameters of the sub-populations introduced in the previous section become themselves random variables, denoted as ***w***, for the frequency, ***µ*** as the mean expression level, and ***v*** for the variance of a sub-population. To avoid confusing notation, in this section we will refer to the mean and variance of the random variables ***w***, ***µ*** and ***v*** solely using the notation 𝔼 [ ] and 𝕍 [ ]. In the general case considered here, these random variables are described by a joint density *h*(*w, µ, v*). Hence, the full population to be studied is constructed from a sample *𝒮* = { (***w***_**1**_, ***µ***_**1**_, ***v***_**1**_), …, (***w***_***i***_, ***µ***_***i***_, ***v***_***i***_) …, (***w***_***N***_**, *µ***_***N***_**, *v***_***N***_)} of *N* vector-valued random variables (***w***, ***µ***, ***v***) sampled from an unknown distribution. A key simplifying assumption made hereafter is that (***w***_***i***_, ***µ***_***i***_, ***v***_***i***_) and (***w***_***j***_, ***µ***_***j***_, ***v***_***j***_) are independent and identically distributed (iid) for all *i ≠ j*. In terms of the frequencies, this is not immediate, since *w*_*i*_ and *w*_*j*_ are dependent due to the constraint of unity sum 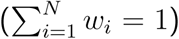. However, this dependency is expected to become negligible, as long as the numbers of cells in each sub-population (see equation 1 of the main text) are iid, and *N* is sufficiently large.

In the following, it is shown that, for a fixed *N*, the mean and variance of the full population are basically “sample estimates” based on 𝒮. Since these estimates are functions of random variables, they are themselves random variables, denoted as _***N***_ ***µ***_***F***_ and _***N***_ ***v***_***F***_, respectively (as in equations 31 and 32):

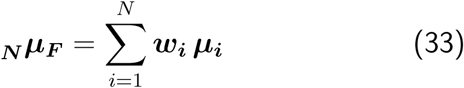

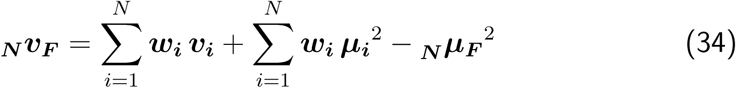

It should be highlighted that ***w***_***i***_, ***µ***_***i***_ and ***v***_***i***_ are, in equations 33 and 34, random variables.

In this framework, one is interested in the expected value of the mean and variance of the full population. Under the law of large numbers, the sample estimates (equations 33 and 34) will converge to the expected values of the mean and variance for sufficiently large *N*. We start by deriving the asymptotic mean of the population:

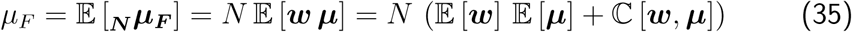

where ℂ [***w***, ***µ***] is the covariance between the random variables ***w*** and ***µ***. Given that 𝔼 [***w***] = 1 /*N*, by definition of the frequencies, one obtains that:

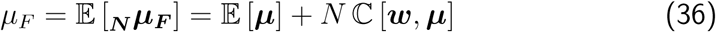

Therefore, it follows that, when the ***w*** and ***µ*** are uncorrelated, the expected mean of the population is simply the expected mean of the sub-populations, corresponding to equation 2 of the main text.

Following a similar reasoning, one obtains the variance:

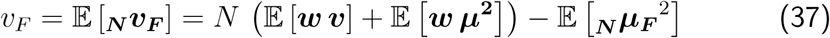

where:

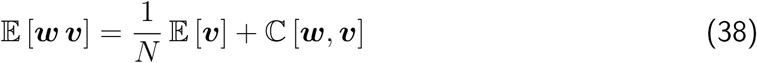

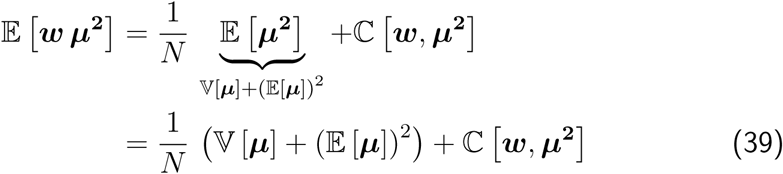

Note the appearance of the term ℂ [***w***, ***µ***^2^], containing the additional random variable ***µ***^2^. Furthermore, the last term in equation 37 can be written as:

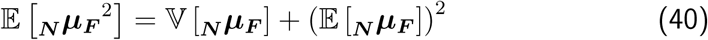

which, using equation 36, becomes:

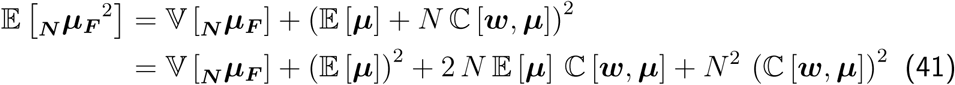

Plugging back equations 38, 39 and 41 into 37, it follows that the variance is given by:

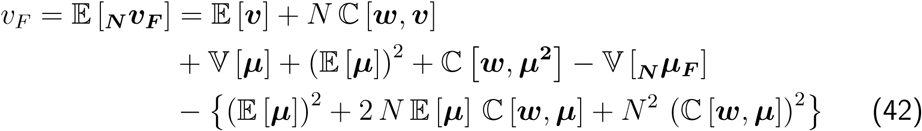

which is reduced to:

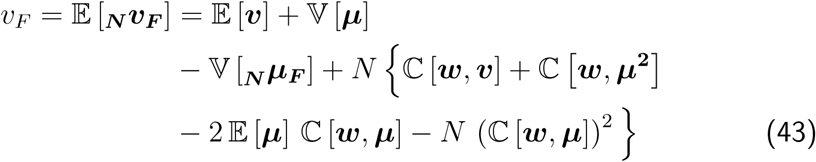

The term 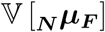 represents an additional contribution, due to variance in the sample mean of the full population as a consequence of sampling, and tends to zero as *N* grows. In this case, provided that there is no correlation between the frequencies (***w***) and either the means (***µ***), the squared means (***µ***^2^) and the variances (***v***) of the sub-population, one obtains equations 2 and 3 of the main text.

### B Basic properties of the logarithmic transformation

In this session, we recall some basic properties of the logarithmic transformation. First of all, recall that, a lognormally-distributed random variable ***x****∼ℒ 𝒩 (µ, σ*) has expected value, variance and coefficient of variation given, respectively, by:

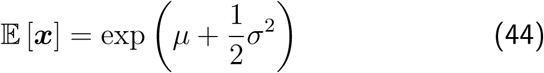

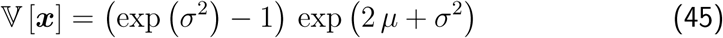

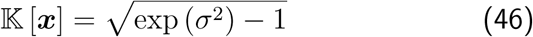

Conversely, the parameters *µ* and *σ* of the lognormal distribution are obtained from 𝔼 [***x***] and 𝕍 [***x***] via:

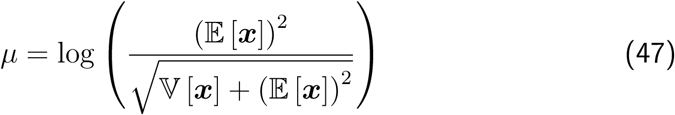

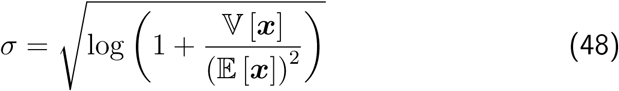

In order to frame the relationship between untransformed and log-transformed values, consider a random variable ***z***, and define ***y*** = log (***z***). If ***y*** can be well approximated by a normal distribution, then equations 44 and 45 can be used to relate the mean and variance of ***z*** and ***y***:

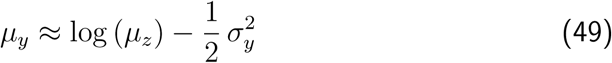

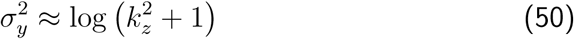

where *k*_*z*_ = 𝕂 [***z***] is the coefficient of variation of ***z***.

### C Model of protein expression in a cell population, for untransformed values

#### C.1 Variation within a sub-population

Starting from equation 4, it follows that a population of cells with dynamics of protein expression levels governed by equations 4 and 5 has stationary mean given by:

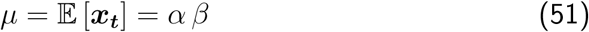

and therefore the stationary mean depends on the average expression rate (therefore, *α*) and on the timescale of protein degradation (*β*). Moreover, the squared stationary coefficient of variation is given by:

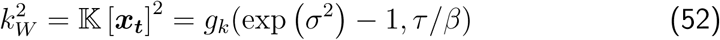

where *g*_*k*_(*·,·*) is an arbitrary function, which can be estimated via simulation (analogous to *g*(*·,·*) in equation 10), and the subscript *W* highlights that the variation is due to the stochastic process influencing the instantaneous rate of protein expression. Hence, the stationary variance is given by:

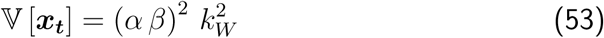

**C.2 Variation among sub-populations**

Following equation 53, the *i*-th sub-population, with parameter *α*_*i*_, has mean and variance of protein levels (see equations 51 and 53):

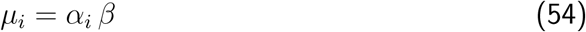

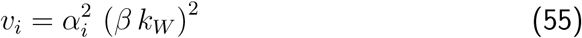

where it should be noted that 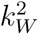 is the same for all sub-populations. Applying equations 2 and 3 of the main text, one obtains that the squared coefficient of variation of the full population is given by:

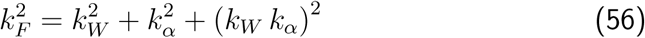

Therefore, equation 56, based on untransformed values, does not follow the simple additive relationship obtained for the variances of log-transformed values (equation 13 of the main text), given the extra term (*k*_*W*_ *k*_*α*_)^2^.

### D Dynamics of the mean of log-transformed values

This section studies the dynamics of the log-transformed mean, to provide a rationale for the exponential-like decay of the function Δ_*H,L*_(*t*) shown in figure 4 (section 4) of the main text based on simulations. The first step is the derivation of a linearised approximation of the log-transformed stochastic model that describes protein expression in a sub-population (defined by equations 4 and 5 of the main text). Afterwards, the dynamics of function Δ_*H,L*_(*t*), which depends on the mean of log-transformed values of high and low expressors, are related to the dynamics of expression levels in the underlying sub-populations.

Since analysis is based on log-transformed values, we define the log-transformed protein level *s*_*t*_:

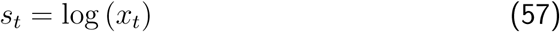

such that:

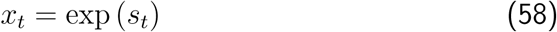

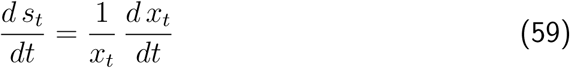

where *x*_*t*_ is the protein level and *y*_*t*_ is the Ornstein-Uhlenbeck process in the original stochastic model (section 2.2.1 of the main text) Therefore, it follows from equation 4 of the main text that:

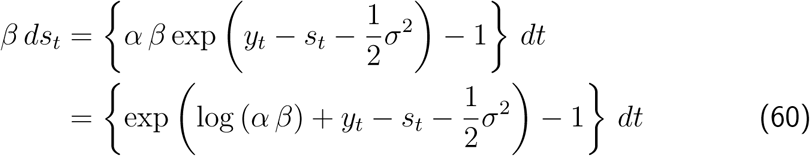

The dynamics of the mean of log-transformed values in a sub-population are then given by:

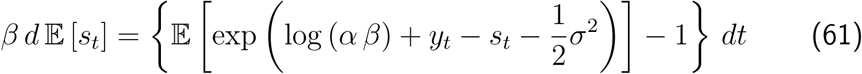

In order to derive an approximation for the function 𝔼 [*s*_*t*_], it is necessary to simplify the term 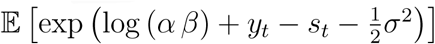. Introducing:

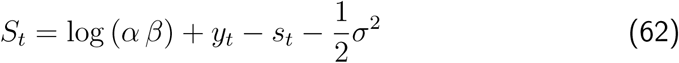

and assuming that it is well-concentrated around a certain instantaneous mean, such that it can be approximated by a normal distribution with mean *m*_*t*_ and variance *v*_*t*_, it follows that (see equation 44):

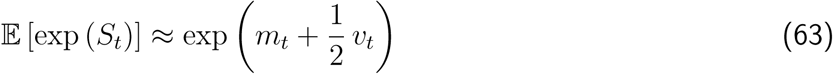

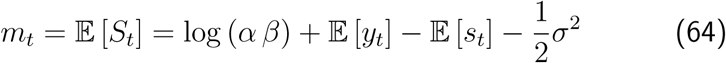

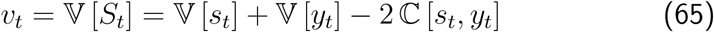

Assuming that 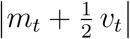 is relatively small, the exponential term in the lefthand side of equation 63 can be linearized:

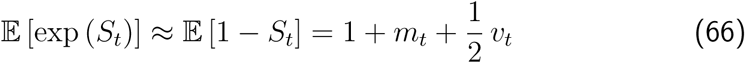

Plugging back into equation 61, one obtains the following linear approximation for the dynamics of the mean of log-transformed values:

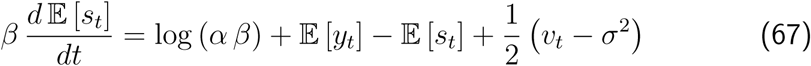

The equation for the mean of the Ornstein-Uhlenbeck process follows as:

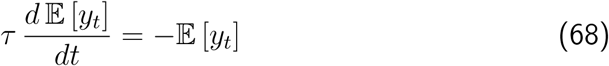

with solution:

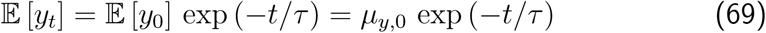

Therefore, one obtains the following equation for *µ*_*t*_ = 𝔼 [*s*_*t*_], which denotes the instantaneous mean of log-transformed values of a single sub-population:

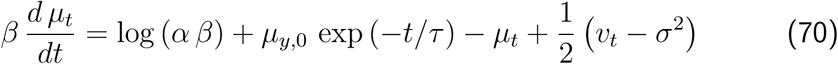

where recall that *v*_*t*_ (equation 65) depends on the variances of log-transformed values of the sub-population (*s*_*t*_) and the Ornstein-Uhlenbeck process variable (*y*_*t*_), besides the covariance between these two.

In terms of the function Δ_*H,L*_(*t*), recall that it is defined as (equation 15 of the main text):

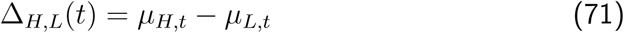

where *µ*_*H,t*_ and *µ*_*L,t*_ are the means of log-transformed values of high and low expressors, respectively, at time *t*. Using equation 36, Δ_*H,L*_(*t*) can be written as:

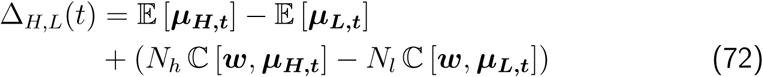

where ***µ***_***H,t***_ and ***µ***_***L,t***_ are random variables denoting the instantaneous mean of a particular sub-population in the high (*N*_*h*_ sub-populations) and low expressors (*N*_*l*_ sub-populations), respectively. Neglecting the term of weighted difference between the covariances in equation 72, it follows that:

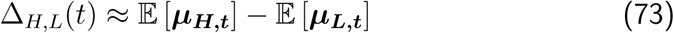

and therefore the dynamics of Δ_*H,L*_(*t*) can be approximated as:

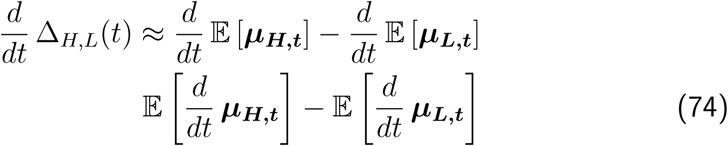

An approximation to the term 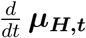 has been derived in equation 70, such that:

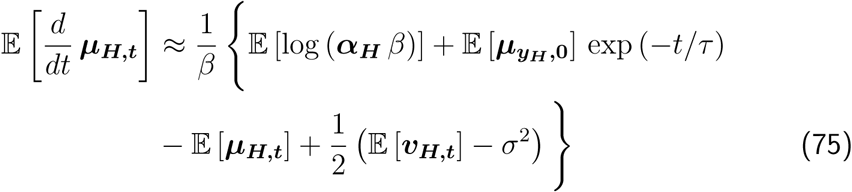

and analogously for the low expressors. Plugging into equation 74, one obtains:

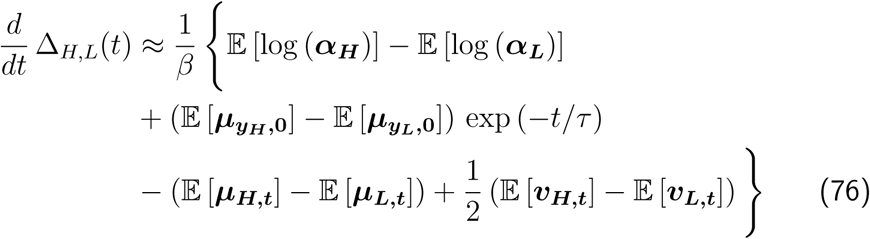

By symmetry, the difference 𝔼 [***v***_***H,t***_] - 𝔼 [***v***_***L,t***_] in equation 76 is expected to be close to zero. Defining the constants:

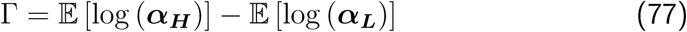

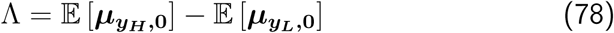

equation 76 is simplified to take the form:

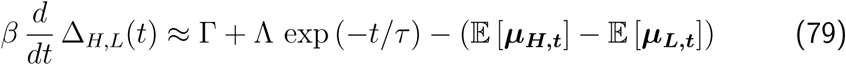

Finally, using the original approximation in equation 73, it follows that:

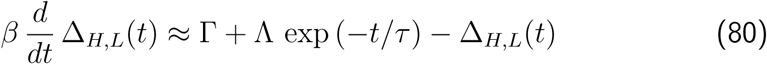

Introducing the auxiliary variable *U (t*) = Λ exp (*-t/τ*), then equation 80 can be written as a two-dimensional linear dynamical system:

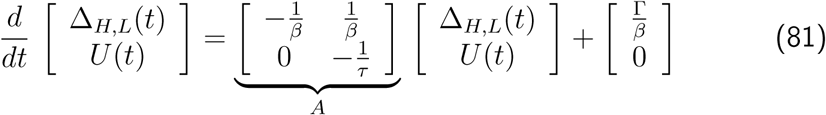

subject to the initial condition:

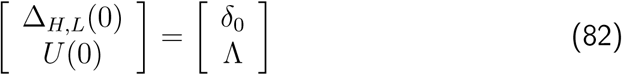

Since the matrix *A* (equation 81) has always non-imaginary eigenvalues, it follows that Δ_*H,L*_(*t*) is the combination of two exponential decays with mean times given by *τ* and *β*, with a single dominant exponential for the extreme cases *β »τ* or *β « τ*.

### E Analysis of the variances of isolated cohorts

In this section, it is shown that analysing the variances of isolated cohorts can provide additional information, given the estimate of 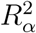. This analysis is based on the same simulation setting considered in section 4. However, it was found that the estimates derived herein are much more sensitive to sampling effects. Therefore, the starting populations considered here had a much larger number of cells (see methods in section E.1).

Figure 10 shows the variance of log-transformed values as a function of time for the “high expressors”. As in section 4, the “all expressors” are also included, as a reference of the starting population. The variance of high expressors is lower than that of the starting population, and either remains constant or increases as a function of time. The same takes place for the low expressors, since the variance is a moment of even order. Finally, as observed for the mean of log values, the asymptotic (stationary) variance is equal to that of the “all expressors” for 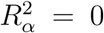, since in this case the unstable component is the only contribution present.

**Figure 10:**
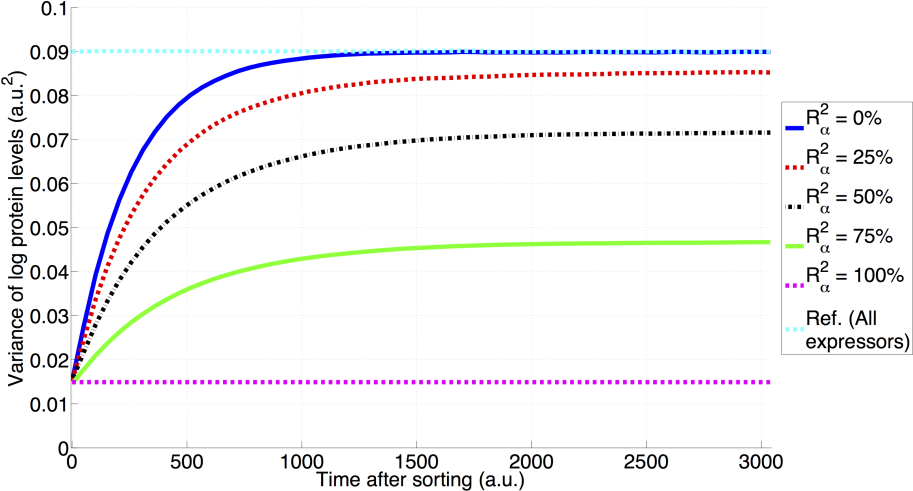
Variance (log values) of “high expressors” after isolation as 10% of starting populations with different values of 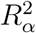, but constant 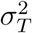. Parameters: *τ* = 500, *β* = 5 and *σ*_*T*_ = 0.3.

Focusing on the asymptotic (stationary) variance, figure 11 shows that the simple, linear, relationship between the mean and 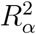 does not hold in this case. In particular, for 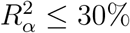, the variance of high and low expressors is very close to that of the all expressors. In order to understand the basis for the relationship showed in figure 11, we consider hereafter the partitioning of the variance of each isolated cohort, in contrast to the main text, which focused on the starting population. However, the value of 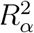 considered will always refer to that of the starting population.

**Figure 11:**
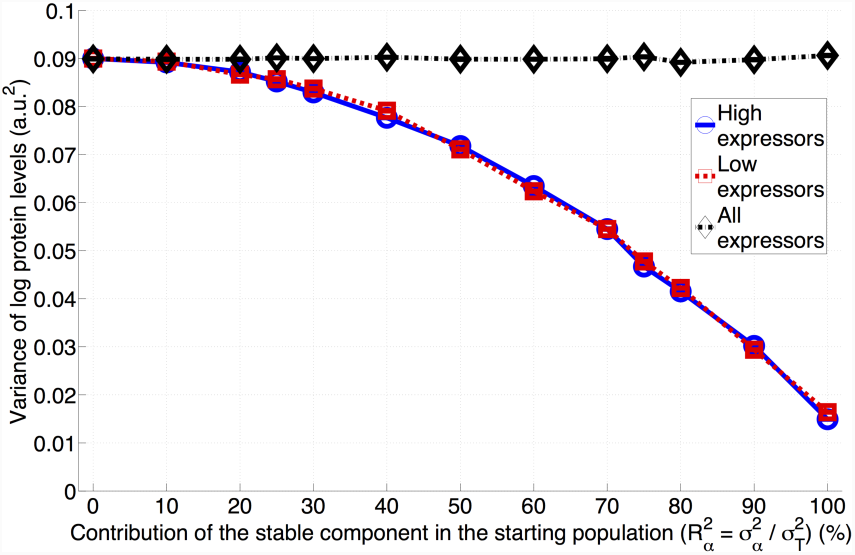
Asymptotic (stationary) variance of expression levels (log values) of high and low expressors, isolated in the simulations as 10% of the starting population, and also of all expressors. The symbols represent the values obtained from the simulations, while the lines represent linear interpolation. Parameters: *τ* = 500, *β* = 5 and *σ*_*T*_ = 0.3.

Recall that the variance of the starting population is given by:

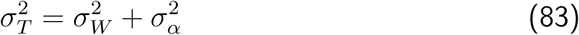

For a general isolated cohort *D*, based on equation 43 (section A of the supplementary information), the variance is partitioned as:

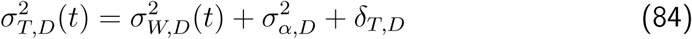

The subscripts *H, L* and *A* in place of *D* will be used to refer to the isolated cohorts corresponding to high, low and all expressors, respectively. The notation 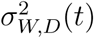 highlights that the variance due to the unstable component becomes a function of time, which will increase until the population becomes stationary. The variance due to the stable component in the isolated cohorts is represented by 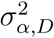, to highlight the fact that it may be different from that of the starting population 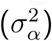 as a consequence of isolating only some sub-populations (see discussion in section 3.2 of the main text). Finally, the term *δ*_*T,D*_ in equation 84 represents a “residual contribution”, which may be introduced by the process of isolating cells. It arises from the covariance terms, which may become nonnegligible even for a starting population satisfying equation 83. Note that, by definition, *δ*_*T,A*_ = 0 (since the “all expressors” satisfy equation 83).

In analogy with the standard F-statistic used for comparing the variances of two samples, we will denote by *F*_*D*_ the ratio between the asymptotic variance of the isolated cohort (equation 83) and the variance of the starting population (84):

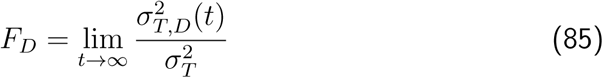

highlighting that this ratio depends only on measurable properties of the two populations. Moreover, define Φ_*D*_ as the ratio between the absolute variances of the stable component in the isolated and starting populations:

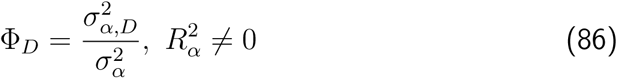

to denote the relative change, as a consequence of isolating cells, in the variance of the stable component in the “new” (isolated) population. In the following, it is shown that *F*_*D*_ and 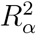 can be used to construct an estimator for Φ_*D*_, denoted as 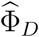. The requirement for 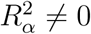 stems from the constraint of 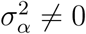.

It follows from taking the ratio between equations 84 and 83 that:

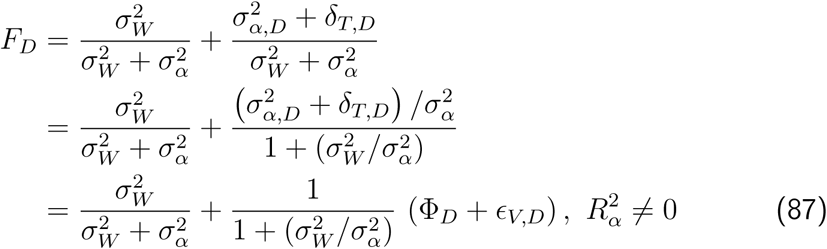

where *ϵ*_*V,D*_ is defined as:

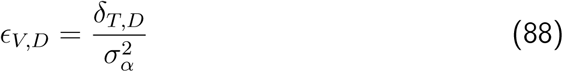

Using the definition of 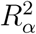:

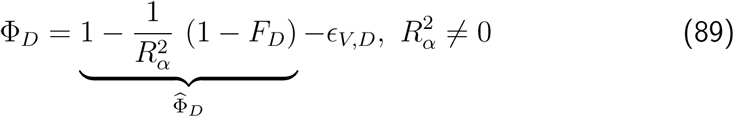

it follows that one estimator for Φ_*D*_, denoted as 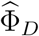, can be obtained via:

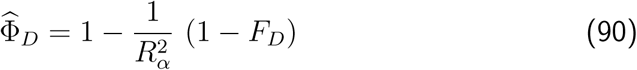

Hence, the “true” and estimated values are related via:

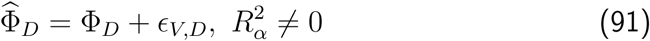

in which *ϵ*_*V,D*_ becomes the bias in the estimation of Φ_*D*_.

Hereafter, we conduct a more detailed analysis of the contributions to the variance in the isolated cohorts, to evaluate the use of the estimator 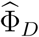 in quantifying Φ_*D*_. This analysis is based on isolating the populations of interest, and simulating until they become stationary. At this point, using the underlying structure of each isolated cohort, the expression levels and number of cells in each sub-population were determined. Using equation 43, the different terms were then calculated.

To understand the basis of the residual contribution (*δ*_*T,D*_), figure 12 depicts the contributions to the asymptotic variance (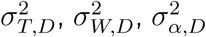 and 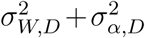) of each of the isolated cohorts, as a function of the value of 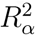. For each isolated cohort, the total variance 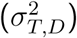 corresponds exactly to the data shown in figure 11, while the term due to the unstable component 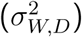, being equal to that in the starting population 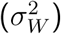, is simply 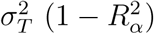. The converse holds for the term arising due to the stable component in the population of all expressors 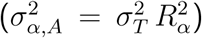, since this population is equivalent to the starting one. On the other hand, for high and low expressors, the value of 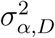, follows a more complicated dependence on 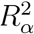, reaching a maximum for values of 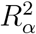 around 60%. Moreover, in these two isolated cohorts, the sum 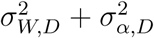 is greater than the total variance 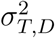, especially for intermediate values of 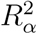. This difference corresponds exactly to the residual component *δ*_*T,D*_. Moreover, as shown in figure 12 (bottom right), it constitutes up to 15% of the total variance, having negative values for all 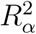. The occurrence of negative values is not unexpected, given that there are both positive and negative terms in equation 43.

**Figure 12:**
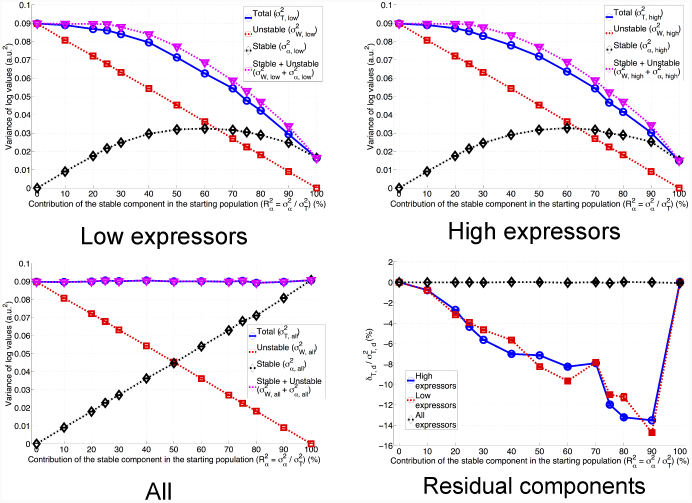
Properties of the isolated cohorts for various values of 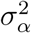, always considering log-transformed expression levels. Shown here are the variance components in the various isolated cohorts, along with the residual component (*δ*_*T,D*_). The latter has been calculated based on equation 43, and the values shown are normalized by the total variance.

However, it is important to recall that the bias *ϵ*_*V,D*_ in the estimation of Φ_*D*_ corresponds to the residual component divided by 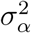. The bias is shown in figure 13 as a function of 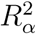, keeping in mind that it is only defined for 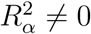. While the residual variance has an absolute value of up to 5% (figure 12, bottom right), it follows that the bias *ϵ*_*V,D*_ varies from −0.18 to 0, vanishing only for 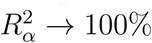. Hence, it is expected that 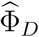 under-estimates Φ_*D*_.

**Figure 13:**
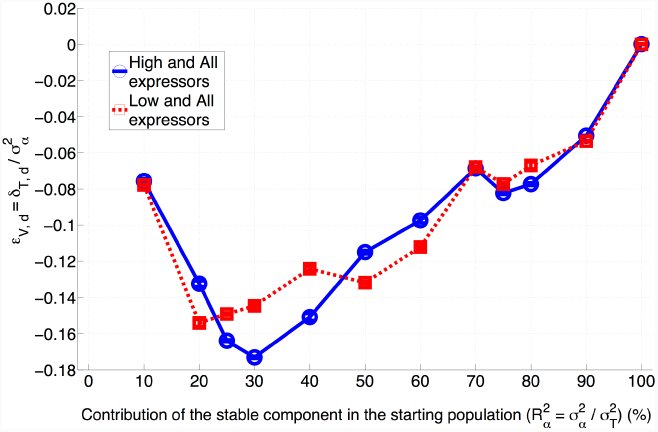
Bias term *ϵ* _*V,D*_ as a function of 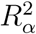 (for 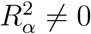), considering the analysis based on the pairs (high, all) and (low, all), determined based on the residual variance (equation 43) and 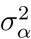.

Finally, the “true” and the estimated values of Φ_*D*_ are shown in figure 14. In this figure, Φ_*D*_ was calculated based on data on the sub-populations composing each isolated cohort, while the estimate 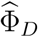 was obtained based on equation 90. These two values are clearly different for values of 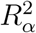 lower than 70%. Figure 14 also shows that, when the bias *ϵ* _*V,D*_ is accounted for (using equation 43, to obtain the residual component and 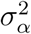), subtracting it from the estimate results in the “true” value. However, given that the bias cannot be estimated in practice, since it depends on the underlying structure of each cell population, it follows that the estimation of Φ_*D*_ via 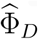 is, indeed, biased in most of the cases.

**Figure 14:**
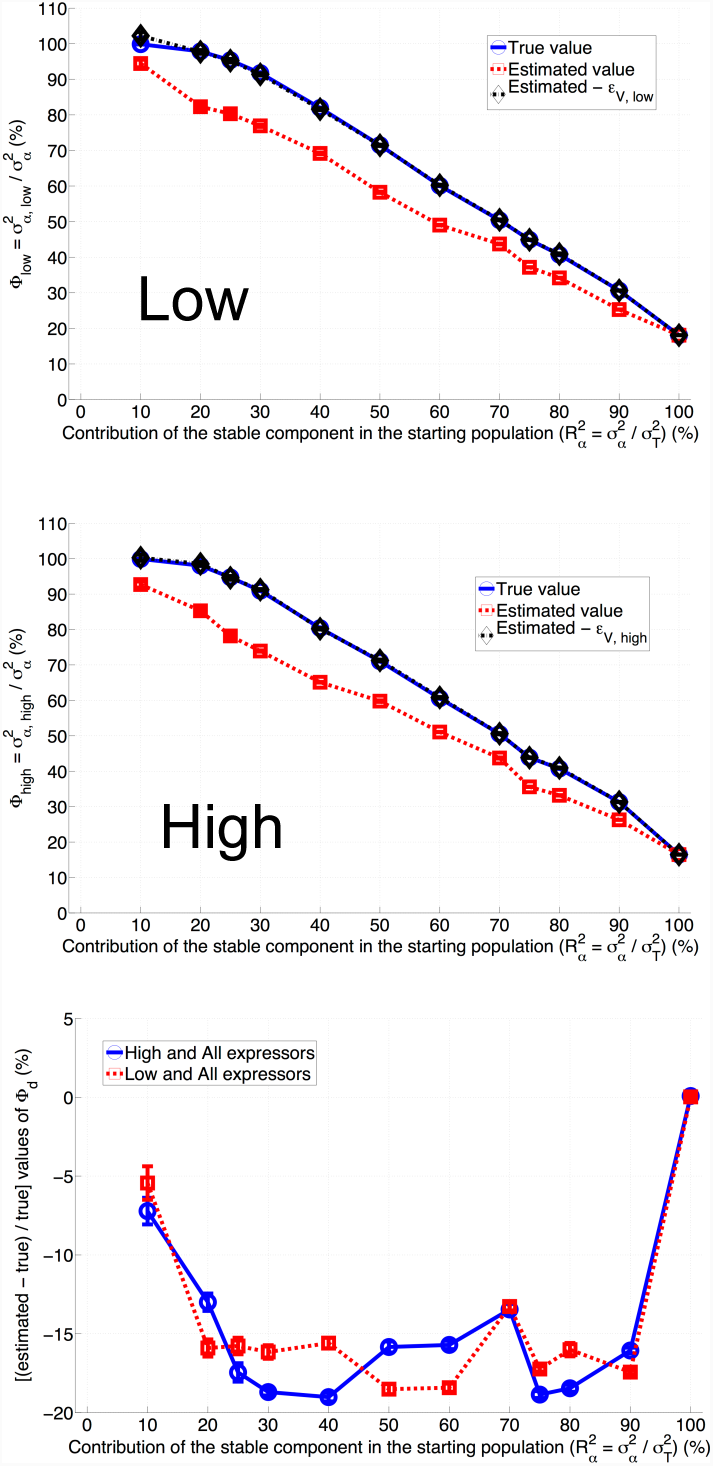
Comparison between the “true” value of Φ_*D*_ and the estimated value 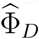 obtained according to equation 90. Each graph also includes the results of subtracting the bias (obtained as in figure 13) from the estimated value, to show that it explains the discrepancy between Φ_*D*_ and 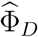.

Therefore, we conclude that the variance can provide additional information. The variance allows to estimate the ratio between the stable component in an isolated cohort (such as the high or low expressors) and the stable component in the starting population (or its proxy the all expressor cohort). This estimate depends on the value of 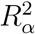, which can be estimated using the approach outlined in section 4 of the main text, and the ratio between the total variances of the two populations being compared (either high and all expressors, or low and all expressors). However, it was shown here that this estimate is biased, due to introduction of a residual component as a consequence of isolating cells based on the expression levels. This bias results typically in an under-estimation of the “true” value of Φ_*D*_ by up to 20% of the true value. Hence, we interpret these results to imply that the asymptotic (stationary) variance is uninformative. Further highlighting the approach based on the means to estimate 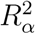, in the case that an estimate of Φ_*D*_ is needed, a simulation-based approach is suggested:

1. estimate 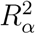 using the analysis of the means
2. using 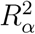, simulate the actual isolation of cells from the starting population, and determine Φ_*D*_

#### E.1 Materials and methods

Simulations were done with the same parameters as those for section 4 of the main text, but with a much larger number of cells in the starting population (30 × 10^6^), and the same number (2 × 10^4^) of sub-populations. Statistics on the sub-populations were calculated based on the asymptotic mean and variance defined by the parameters 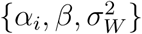 describing each sub-population.

### F Forward scatter distributions in sorted cohorts of high and low expressors

**Figure 15:**
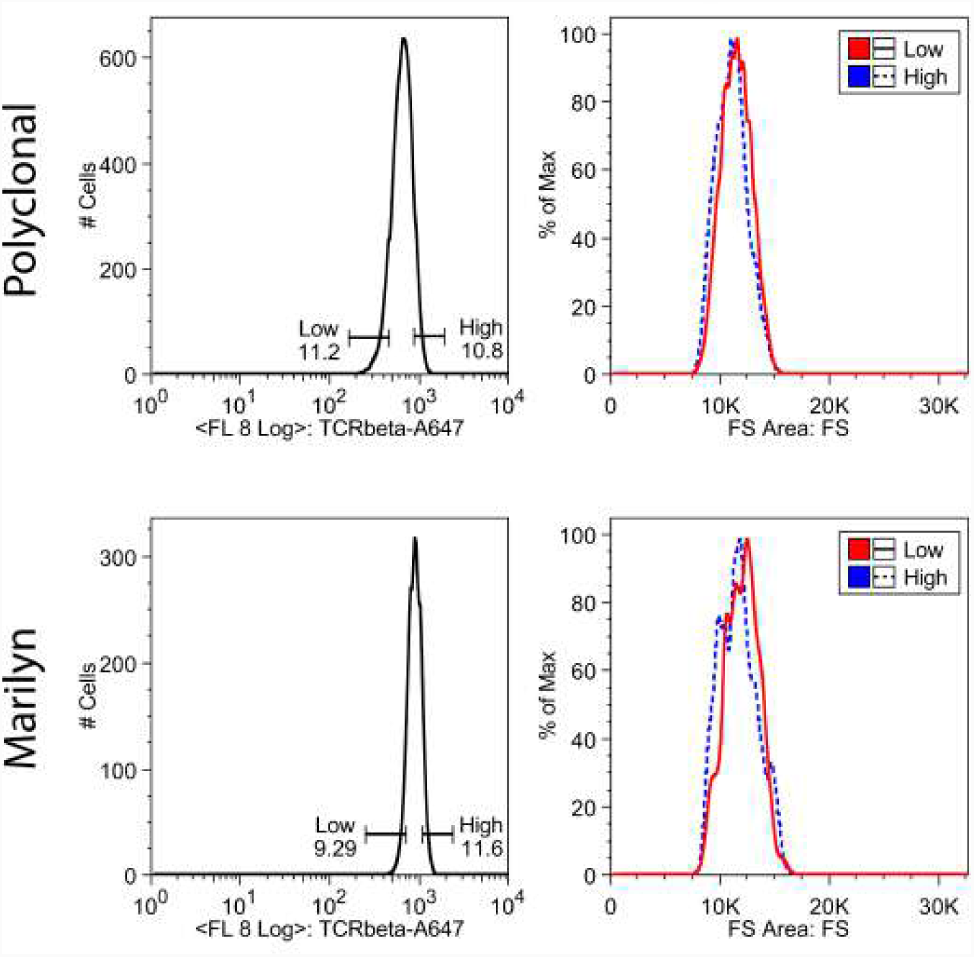
Forward scatter distributions in cohorts of high and low TCR expressors sorted from naive polyclonal (top) and Marilyn monoclonal (bottom) CD4 T cells. Left: Histograms of TCR intensity in the cell populations are shown on the left together with the gates used to sorting the high and low expressor cohorts. The numbers are the percentage of cells in each cohort. Right: Histograms of Foreward Scatter Area for the high (blue) and low (red) expressors immediately after sorting.

## References

[1] Matthew V Rockman and Leonid Kruglyak. Genetics of global gene expression. Nature Reviews Genetics, 7(11):862–72, November 2006. ISSN 1471-0056. URL http://www.ncbi.nlm.nih.gov/pubmed/17047685.

[2] Frank W. Albert and Leonid Kruglyak. The role of regulatory variation in complex traits and disease. Nature Reviews Genetics, 16:197 EP –, Feb 24, 2015. URL https://doi.org/10.1038/nrg3891. Review Article.

[3] What is your conceptual definition of “cell type” in the context of a mature organism? Cell Systems, 4(3):255–259, mar 2017. URL https://doi.org/10.1016/j.cels.2017.03.006.

[4] Michael B Elowitz, Arnold J Levine, Eric D Siggia, and Peter S Swain. Stochastic gene expression in a single cell. Science (New York, N.Y.), 297 (5584):1183–6, August 2002. ISSN 1095-9203. URL http://www.ncbi.nlm.nih.gov/pubmed/12183631.

[5] Jonathan M Raser and Erin K O’Shea. Control of stochasticity in eukaryotic gene expression. Science (New York, N.Y.), 304(5678):1811–4, June 2004. ISSN 1095-9203. URL http://www.pubmedcentral.nih.gov/articlerender.fcgi?artid=1410811\&tool=pmcentrez\&rendertype=abstract.

[6] Juan M Pedraza and Alexander van Oudenaarden. Noise propagation in gene networks. Science (New York, N.Y.), 307(5717):1965–9, March 2005. ISSN 1095-9203. URL http://www.ncbi.nlm.nih.gov/pubmed/15790857.

[7] Alex Sigal, Ron Milo, Ariel Cohen, Naama Geva-Zatorsky, Yael Klein, Yu-valal Liron, Nitzan Rosenfeld, Tamar Danon, Natalie Perzov, and Uri Alon. Variability and memory of protein levels in human cells. Nature, 444(7119): 643–6, November 2006. ISSN 1476-4687. URL http://www.ncbi.nlm.nih.gov/pubmed/17122776.

[8] Arjun Raj, Scott a Rifkin, Erik Andersen, and Alexander van Oudenaarden. Variability in gene expression underlies incomplete penetrance. Nature, 463(7283):913–8, March 2010. ISSN 1476-4687. URL http://www.pubmedcentral.nih.gov/articlerender.fcgi?artid=2836165\&tool=pmcentrez\&rendertype=abstract.

[9] Arjun Raj and Alexander van Oudenaarden. Nature, nurture, or chance: stochastic gene expression and its consequences. Cell, 135(2):216–26, October 2008. ISSN 1097-4172. URL http://www.pubmedcentral.nih.gov/articlerender.fcgi?artid=3118044\&tool=pmcentrez\&rendertype=abstract.

[10] Sui Huang. Non-genetic heterogeneity of cells in development: more than just noise. Development (Cambridge, England), 136(23):3853–62, December 2009. ISSN 1477-9129. URL http://www.pubmedcentral.nih.gov/articlerender.fcgi?artid=2778736\&tool=pmcentrez\&rendertype=abstract.

[11] Qiaolin Deng, Daniel Ramsköld, Björn Reinius, and Rickard Sandberg. Single-cell rna-seq reveals dynamic, random monoallelic gene expression in mammalian cells. Science, 343(6167):193–196, 2014. ISSN 0036-8075. URL http://science.sciencemag.org/content/343/6167/193.

[12] Roy D Dar, Brandon S Razooky, Abhyudai Singh, Thomas V Trimeloni, James M. McCollum, Chris D. Cox, Michael L. Simpson, and Leor S. Weinberger. Transcriptional burst frequency and burst size are equally modulated across the human genome. Proceedings of the National Academy of Sciences of the United States of America, 109(43):17454–9, October 2012. ISSN 1091-6490. URL http://www.pnas.org/cgi/doi/10.1073/pnas.1213530109 http://www.ncbi.nlm.nih.gov/pubmed/23064634.

[13] Tiago Paixão, Tiago P Carvalho, Dinis P Calado, and Jorge Carneiro. Quantitative insights into stochastic monoallelic expression of cytokine genes. Immunology and Cell Biology, 85(4):315–22, June 2007. ISSN 0818-9641. URL http://www.ncbi.nlm.nih.gov/pubmed/17438562.

[14] Tibor Kalmar, Chea Lim, Penelope Hayward, Silvia Munõz Descalzo, Jennifer Nichols, Jordi Garcia-Ojalvo, and Alfonso Martinez Arias. Regulated fluctuations in nanog expression mediate cell fate decisions in embryonic stem cells. PLoS Biology, 7(7):e1000149, July 2009. ISSN 1545-7885. URL http://www.pubmedcentral.nih.gov/articlerender.fcgi?artid=2700273\&tool=pmcentrez\&rendertype=abstract.

[15] Luca Mariani, Edda G Schulz, Maria H Lexberg, Caroline Helmstetter, Andreas Radbruch, Max Löhning, and Thomas Höfer. Short-term memory in gene induction reveals the regulatory principle behind stochastic IL-4 expression. Molecular Systems Biology, 6(359):359, April 2010. ISSN 1744-4292. URL http://www.pubmedcentral.nih.gov/articlerender.fcgi?artid=2872609\&tool=pmcentrez\&rendertype=abstract.

[16] Hannah H Chang, Martin Hemberg, Mauricio Barahona, Donald E Ingber, and Sui Huang. Transcriptome-wide noise controls lineage choice in mammalian progenitor cells. Nature, 453(7194):544–7, May 2008. ISSN 1476-4687. URL http://www.ncbi.nlm.nih.gov/pubmed/18497826.

[17] Cristina Pina, Cristina Fugazza, Alex J Tipping, John Brown, Shamit Soneji, Jose Teles, Carsten Peterson, and Tariq Enver. Inferring rules of lineage commitment in haematopoiesis. Nature Cell Biology, 14(3):287–94, January 2012. ISSN 1476-4679. URL http://www.ncbi.nlm.nih.gov/pubmed/ 22344032.

[18] Daniel R. Sisan, Michael Halter, Joseph B. Hubbard, and Anne L. Plant. Predicting rates of cell state change caused by stochastic fluctuations using a data-driven landscape model. Proceedings of the National Academy of Sciences of the United States of America, 109(47):19262–7, November 2012. ISSN 1091-6490. URL http://www.pnas.org/cgi/doi/10.1073/pnas.1207544109 http://www.ncbi.nlm.nih.gov/pubmed/23115330 http://www.pubmedcentral.nih.gov/articlerender.fcgi?artid=3511108\&tool=pmcentrez\&rendertype=abstract.

[19] Peter S Swain, Michael B Elowitz, and Eric D Siggia. Intrinsic and extrinsic contributions to stochasticity in gene expression. Proceedings of the National Academy of Sciences of the United States of America, 99(20):12795–800, October 2002. ISSN 0027-8424. URL http://www.pubmedcentral.nih.gov/articlerender.fcgi?artid=130539\&tool=pmcentrez\&rendertype=abstract.

[20] Mary J Dunlop, Robert Sidney Cox, Joseph H Levine, Richard M Murray, and Michael B Elowitz. Regulatory activity revealed by dynamic correlations in gene expression noise. Nature genetics, 40(12):1493–8, December 2008. ISSN 1546-1718. URL http://www.pubmedcentral.nih.gov/articlerender.fcgi?artid=2829635\&tool=pmcentrez\&rendertype=abstract.

[21] Julia Rausenberger and Markus Kollmann. Quantifying origins of cell-to-cell variations in gene expression. Biophysical Journal, 95(10):4523–8, November 2008. ISSN 1542-0086. URL http://www.pubmedcentral.nih.gov/articlerender.fcgi?artid=2576406\&tool=pmcentrez\&rendertype=abstract.

[22] Brian Munsky, Brooke Trinh, and Mustafa Khammash. Listening to the noise: random fluctuations reveal gene network parameters. Molecular systems biology, 5(318):318, January 2009. ISSN 1744-4292. URL http://www.pubmedcentral.nih.gov/articlerender.fcgi?artid=2779089\&tool=pmcentrez\&rendertype=abstract.

[23] Ruty Rinott, Ariel Jaimovich, and Nir Friedman. Exploring transcription regulation through cell-to-cell variability. Proceedings of the National Academy of Sciences of the United States of America, 108(15):6329–34, April 2011. ISSN 1091-6490. URL http://www.pubmedcentral.nih.gov/articlerender.fcgi?artid=3076844\&tool=pmcentrez\&rendertype=abstract.

[24] Michal Komorowski, Jacek Miekisz, and Michael P.H. Stumpf. Decomposing Noise in Biochemical Signaling Systems Highlights the Role of Protein Degradation. Biophysical Journal, 104(8):1783–1793, April 2013. ISSN 00063495. URL http://linkinghub.elsevier.com/retrieve/pii/S0006349513002452.

[25] S H Orkin. Diversification of haematopoietic stem cells to specific lineages. Nature reviews. Genetics, 1(1):57–64, October 2000. ISSN 1471-0056. URL http://www.ncbi.nlm.nih.gov/pubmed/11262875.

[26] Myriam Hemberger, Wendy Dean, and Wolf Reik. Epigenetic dynamics of stem cells and cell lineage commitment: digging Waddington’s canal. Nature reviews. Molecular cell biology, 10(8):526–37, August 2009. ISSN 1471-0080. URL http://www.ncbi.nlm.nih.gov/pubmed/19603040.

[27] O Lantz, I Grandjean, P Matzinger, and J P Di Santo. Gamma chain required for naïve CD4+ T cell survival but not for antigen proliferation. Nature Immunology, 1(1):54–8, July 2000. ISSN 1529-2908. URL http://www.ncbi.nlm.nih.gov/pubmed/10881175.

[28] Daniel Gianola, Bjorg Heringstad, and Jorgen Odegaard. On the quantitative genetics of mixture characters. Genetics, 173(4):2247–55, August 2006. ISSN 0016-6731. URL http://www.pubmedcentral.nih.gov/articlerender.fcgi?artid=1569724\&tool=pmcentrez\&rendertype=abstract.

[29] Vahid Shahrezaei, Julien F Ollivier, and Peter S Swain. Colored extrinsic fluctuations and stochastic gene expression. Molecular Systems Biology, 4(196):196, January 2008. ISSN 1744-4292. URL http://www.pubmedcentral.nih.gov/articlerender.fcgi?artid=2424296\&tool=pmcentrez\&rendertype=abstract.

[30] Ludwig Arnold. Stochastic differential equations: theory and applications. Wiley, 1 edition, 1974. ISBN 0471033596.

[31] Abhyudai Singh, Brandon S Razooky, Roy D Dar, and Leor S Weinberger. Dynamics of protein noise can distinguish between alternate sources of geneexpression variability. Molecular Systems Biology, 8(607):1–9, August 2012. ISSN 1744-4292. URL http://www.nature.com/doifinder/10.1038/msb.2012.38.

[32] Nitzan Rosenfeld, Jonathan W Young, Uri Alon, Peter S Swain, and Michael B Elowitz. Gene regulation at the single-cell level. Science (New York, N.Y.), 307(5717):1962–5, March 2005. ISSN 1095-9203. URL http://www.ncbi.nlm.nih.gov/pubmed/15790856.

[33] Gerald P Morris and Paul M Allen. How the TCR balances sensitivity and specificity for the recognition of self and pathogens. Nature Immunology, 13(2):121–8, February 2012. ISSN 1529-2916. URL http://www.ncbi.nlm.nih.gov/pubmed/22261968.

[34] Ute Koch and Freddy Radtke. Mechanisms of T cell development and transformation. Annual Review of Cell and Developmental Biology, 27: 539–62, January 2011. ISSN 1530-8995. URL http://www.ncbi.nlm.nih.gov/pubmed/21740230.

[35] Isabelle Grandjean, Livine Duban, Elizabeth a Bonney, Erwan Corcuff, James P Di Santo, Polly Matzinger, and Olivier Lantz. Are major histocompatibility complex molecules involved in the survival of naive CD4+ T cells? The Journal of Experimental Medicine, 198(7):1089–102, October 2003. ISSN 0022-1007. URL http://www.pubmedcentral.nih.gov/articlerender.fcgi?artid=2194222\&tool=pmcentrez\&rendertype=abstract.

[36] Fatima-Zahra El Hentati, Frederic Gruy, Cristina Iobagiu, and Claude Lam-ert. Variability of cd3 membrane expression and t cell activation capacity. Cytometry Part B: Clinical Cytometry, 78B(2):105–114, 2010. URL https://onlinelibrary.wiley.com/doi/abs/10.1002/cyto.b.20496.

[37] Brenna L Brady, Natalie C Steinel, and Craig H Bassing. Antigen receptor allelic exclusion: an update and reappraisal. Journal of Immunology (Baltimore, Md.: 1950), 185(7):3801–8, October 2010. ISSN 1550-6606. URL http://www.pubmedcentral.nih.gov/articlerender.fcgi?artid=3008371\&tool=pmcentrez\&rendertype=abstract.

[38] Elissa K Deenick, Amanda V Gett, and Philip D Hodgkin. Stochastic model of T cell proliferation: a calculus revealing IL-2 regulation of precursor frequencies, cell cycle time, and survival. Journal of Immunology (Baltimore, Md.: 1950), 170(10):4963–72, May 2003. ISSN 0022-1767. URL http://www.ncbi.nlm.nih.gov/pubmed/12734339.

[39] W T Lee, X M Yin, and E S Vitetta. Functional and ontogenetic analysis of murine CD45Rhi and CD45Rlo CD4+ T cells. Journal of Immunology (Baltimore, Md.: 1950), 144(9):3288–95, May 1990. ISSN 0022-1767. URL http://www.ncbi.nlm.nih.gov/pubmed/1970350.

[40] Kenneth P Burnham and David R Anderson. Model Selection and Inference: A Practical Information-Theoretic Approach. Springer-Verlag, New York, NY, 1998. ISBN 0-387-98504-2.

[41] Frank Johannes, Vincent Colot, and Ritsert C Jansen. Epigenome dynamics: a quantitative genetics perspective. Nature Reviews Genetics, 9(11):883–90, November 2008. ISSN 1471-0064. URL http://www.ncbi.nlm.nih.gov/pubmed/18927581.

[42] Warren Pilbrough, Trent P. Munro, and Peter Gray. Intraclonal protein expression heterogeneity in recombinant cho cells. PLOS ONE, 4(12):1–11, 12 2009. URL https://doi.org/10.1371/journal.pone.0008432.

[43] Y. Taniguchi, P. J. Choi, G.-W. Li, H. Chen, M. Babu, J. Hearn, A. Emili, and X. S. Xie. Quantifying e. coli proteome and transcriptome with single-molecule sensitivity in single cells. Science, 329(5991):533–538, jul 2010. URL https://doi.org/10.1126/science.1188308.

[44] David Redmond, Asaf Poran, and Olivier Elemento. Single-cell tcrseq: paired recovery of entire t-cell alpha and beta chain transcripts in t-cell receptors from single-cell rnaseq. Genome Medicine, 8(1):80, Jul 27, 2016. ISSN 1756-994X. URL https://doi.org/10.1186/s13073-016-0335-7.

[45] Sabrina L Spencer and Peter K Sorger. Measuring and modeling apoptosis in single cells. Cell, 144(6):926–39, March 2011. ISSN 1097-4172. URL http://www.pubmedcentral.nih.gov/articlerender.fcgi?artid=3087303\&tool=pmcentrez\&rendertype=abstract.

[46] K Smith, B Seddon, M aPurbhoo, R Zamoyska, a G Fisher, and M Merkenschlager. Sensory adaptation in naive peripheral CD4 T cells. The Journal of Experimental Medicine, 194(9):1253–61, November 2001. ISSN 0022-1007. URL http://www.pubmedcentral.nih.gov/articlerender.fcgi?artid=2195983\&tool=pmcentrez\&rendertype=abstract.

[47] Megan J Palmer, Vinay S Mahajan, Jianzhu Chen, Darrell J Irvine, and Douglasa Lauffenburger. Signaling thresholds govern heterogeneity in IL-7-receptor-mediated responses of naïve CD8(+) T cells. Immunology and Cell Biology, 89(5):581–94, July 2011. ISSN 1440-1711. URL http://www.ncbi.nlm.nih.gov/pubmed/21339767.

[48] Judith N. Mandl, João P. Monteiro, Nienke Vrisekoop, and Ronald N. Germain. T cell-positive selection uses self-ligand binding strength to optimize repertoire recognition of foreign antigens. Immunity, 38(2): 263–74, February 2013. ISSN 1097-4180. URL http://linkinghub.elsevier.com/retrieve/pii/S1074761312004608 http://www.ncbi.nlm.nih.gov/pubmed/23290521.

[49] Charles Sinclair, Manoj Saini, Ina Schim van der Loeff, Shimon Sakaguchi, and Benedict Seddon. The Long-Term Survival Potential of Mature T Lym-phocytes Is Programmed During Development in the Thymus. Science Sig-naling, 4(199):ra77, January 2011. ISSN 1937-9145. URL http://www.ncbi.nlm.nih.gov/pubmed/22087033.

[50] Thiago Guzella. variation in protein expression levels in cell populations. PhD thesis, Instituto de Tecnologia Quimica e Biologica, Universidade Nova de Lisboa, 2013.

[51] Samuel Marguerat, Alexander Schmidt, Sandra Codlin, Wei Chen, Ruedi Aebersold, and Jurg Bahler. Quantitative analysis of fission yeast transcrip-tomes and proteomes in proliferating and quiescent cells. Cell, 151(3):671–683, 2012. ISSN 0092-8674. URL http://www.sciencedirect.com/science/article/pii/S0092867412011269.

[52] Jose M Polo, Susanna Liu, Maria Eugenia Figueroa, Warakorn Kulalert, Sarah Eminli, Kah Yong Tan, Effie Apostolou, Matthias Stadtfeld, Yushan Li, Toshi Shioda, Sridaran Natesan, Amy J Wagers, Ari Melnick, Todd Evans, and Konrad Hochedlinger. Cell type of origin influences the molecular and functional properties of mouse induced pluripotent stem cells. Nature Biotechnology, 28(8):848–55, August 2010. ISSN 1546-1696. URL http://www.pubmedcentral.nih.gov/articlerender.fcgi?artid=3148605\&tool=pmcentrez\&rendertype=abstract.

[53] Jacob Hanna, Krishanu Saha, Bernardo Pando, Jeroen van Zon, Christo-pher J Lengner, Menno P Creyghton, Alexander van Oudenaarden, and Rudolf Jaenisch. Direct cell reprogramming is a stochastic process amenable to acceleration. Nature, 462(7273):595–601, December 2009. ISSN 1476-4687. URL http://www.pubmedcentral.nih.gov/articlerender.fcgi?artid=2789972\&tool=pmcentrez\&rendertype=abstract.

[54] Philipp Thomas. Making sense of snapshot data: ergodic principle for clonal cell populations. Journal of The Royal Society Interface, 14(136), 2017. ISSN 1742-5689. URL http://rsif.royalsocietypublishing.org/content/14/136/20170467.

[55] H Liu, M Rhodes, D L Wiest, and D a Vignali. On the dynamics of TCR:CD3 complex cell surface expression and downmodulation. Immunity, 13(5):665–75, November 2000. ISSN 1074-7613. URL http://www.ncbi.nlm.nih.gov/pubmed/11114379.

[56] N Bronstein-Sitton, Lynn Wang, Leonor Cohen, and Michal Baniyash. Expression of the T cell antigen receptor zeta chain following activation is controlled at distinct checkpoints. Implications for cell surface receptor downmodulation and re-expression. The Journal of Biological Chemistry, 274 (33):23659–65, August 1999. ISSN 0021-9258. URL http://www.ncbi.nlm.nih.gov/pubmed/10438549.

[57] J Sousa and J Carneiro. A mathematical analysis of TCR serial triggering and down-regulation. European Journal of Immunology, 30(11):3219–27, November 2000. ISSN 0014-2980. URL http://www.ncbi.nlm.nih.gov/pubmed/11093137.

[58] Ian B Dodd, Mille a Micheelsen, Kim Sneppen, and Geneviéve Thon. Theoretical analysis of epigenetic cell memory by nucleosome modification. Cell, 129(4):813–22, May 2007. ISSN 0092-8674. URL http://www.ncbi.nlm.nih.gov/pubmed/17512413.

[59] Marzena Gajecka. Unrevealed mosaicism in the next-generation sequencing era. Molecular Genetics and Genomics, 291(2):513–530, Apr 01, 2016. ISSN 1617-4623. URL https://doi.org/10.1007/s00438-015-1130-7.

[60] William G Hill. Understanding and using quantitative genetic variation. Philosophical Transactions of the Royal Society of London. Series B, Biological Sciences, 365(1537):73–85, January 2010. ISSN 1471-2970. URL http://www.pubmedcentral.nih.gov/articlerender.fcgi?artid=2842708\&tool=pmcentrez\&rendertype=abstract.

[61] Andreas Hilfinger and Johan Paulsson. Separating intrinsic from extrinsic fluctuations in dynamic biological systems. Proceedings of the National Academy of Sciences of the United States of America, 108(29):12167–72, July 2011. ISSN 1091-6490. URL http://www.pubmedcentral.nih.gov/articlerender.fcgi?artid=3141918\&tool=pmcentrez\&rendertype=abstract.

[62] Ben D MacArthur, Ana Sevilla, Michel Lenz, Franz-Josef Müller, Berhard M Schuldt, Andreas a Schuppert, Sonya J Ridden, Patrick S Stumpf, Miguel Fidalgo, Avi Ma’ayan, Jianlong Wang, and Ihor R Lemischka. Nanog-dependent feedback loops regulate murine embryonic stem cell heterogeneity. Nature Cell Biology, 14(11):1–11, October 2012. ISSN 1476-4679. URL http://www.ncbi.nlm.nih.gov/pubmed/23103910.

[63] Fahim Farzadfard and Timothy K. Lu. Emerging applications for DNA writers and molecular recorders. Science, 361(6405):870–875, aug 2018. URL https://doi.org/10.1126/science.aat9249.

[64] Y Ghendler, Alex Smolyar, H C Chang, and Ellis L Reinherz. One of the CD3epsilon subunits within a T cell receptor complex lies in close proximity to the Cbeta FG loop. The Journal of Experimental Medicine, 187(9):1529–36, May 1998. ISSN 0022-1007. URL http://www.pubmedcentral.nih.gov/articlerender.fcgi?artid=2212265\&tool=pmcentrez\&rendertype=abstract.

